# Epigenome erosion in Alzheimer’s disease brain cells and induced neurons

**DOI:** 10.1101/2023.10.15.562394

**Authors:** Bang-An Wang, Jeffrey R. Jones, Jingtian Zhou, Wei Tian, Yue Wu, Wenliang Wang, Peter Berube, Anna Bartlett, Rosa Castanon, Joseph R. Nery, Huaming Chen, Mia Kenworthy, Jordan Altshul, Cynthia Valadon, Yichen Wang, Austin Kang, Ryan Goodman, Michelle Liem, Naomi Claffey, Caz O’Connor, Jeffrey Metcalf, Chongyuan Luo, Fred H. Gage, Joseph R. Ecker

## Abstract

Late-onset Alzheimer’s disease (LOAD) is typically sporadic, correlated only to advanced age, and has no clear genetic risk factors. The sporadic nature of LOAD presents a challenge to understanding its pathogenesis and mechanisms. Here, we comprehensively investigated the epigenome of LOAD primary entorhinal cortex brain tissues via single-cell multi-omics technologies, simultaneously capturing DNA methylation and 3D chromatin conformation. We identified AD-specific DNA methylation signatures and found they interact with bivalent promoters of AD differentially expressed genes. In addition, we discovered global chromosomal epigenome erosion of 3D genome structure within and across brain cell types. Furthermore, to evaluate whether these age- and disease-dependent molecular signatures could be detected in the *in vitro* cellular models, we derived induced neurons (iNs) converted directly from AD patients’ fibroblasts and found a set of conserved methylation signatures and shared molecular processes. We developed a machine-learning algorithm to identify robust and consistent methylation signatures of LOAD *in vivo* primary brain tissues and *in vitro* fibroblast-derived iNs. The results recapitulate the age- and disease-related epigenetic features in iNs and highlight the power of epigenome and chromatin conformation for identifying molecular mechanisms of neuronal aging and generating biomarkers for LOAD.

**HIGHLIGHT:**

1. AD-specific DNA methylation signatures are identified in entorhinal cortex brain cell types
2. The AD differentially expressed genes linked with differentially methylated regions via loop interactions are enriched in a bivalent chromatin state
3. Chromosomal epigenome erosion of 3D genome structures occurs in LOAD brain cell types.
4. Shared and reliable methylation signatures are observed in both *in vitro* cellular iN models and primary brain tissues.
5. Machine learning models identify robust and reliable methylation loci as AD biomarkers across cell types.

## INTRODUCTION

Studying human age-dependent disorders is a long-standing challenge, especially for inaccessible tissues like the human brain. Sporadic late-onset Alzheimer’s disease (LOAD) accounts for 95% of all AD cases.^1^ Unlike the early-onset familial AD that is linked to genetic mutations in a specific gene, such as those found in APP, PSEN1 and PSEN2 genes, LOAD is thought to be caused by a complex combination of multiple genes and environmental factors, largely aligning to several age-related co-morbidities. Elucidating the complex genetic background interactions and epigenetic regulation that likely contribute to LOAD is critical to developing targeted therapies.^2^ DNA methylation, the most studied epigenetic system in mammals, has been confirmed to play a crucial role in multiple human diseases such as cancer, imprinting and repeat-instability disorders.^3^ Intriguingly, aberrant DNA methylation is observed in normal aging processes, highlighting the link between proper epigenetic regulation and age-dependent cellular functions.

To characterize genome-wide LOAD-specific methylation signatures from *in vivo* brain cell types, aligning our work with current brain cell type atlas efforts led by BRAIN Initiative Cell Census Network (BICCN),^4–6^ we performed single-nucleus methyl-3C sequencing (sn-m3C-seq), to jointly profile chromatin conformation and methylome from the same cell.^7,8^ This approach enabled the definition of the cell type taxonomy in AD patients and identified differentially methylated regions between AD and control (aDMRs) within and across brain cell types and revealing erosion of the epigenome in single brain cells of LOAD patients based on cell type-specific 3D genome structure alterations. These findings in human AD patients are consistent with the observations on loss of epigenetic information in aged mice^9^ and more recently in an AD mouse model^10^ and human aged cerebellar granule neurons.^11^

In addition, to assess whether the epigenetic signatures found from *in vivo* human brain tissues can be detected in cellular models, we derived induced neurons (iNs) directly converted from dermal fibroblasts of LOAD patients and generated snmCT-seq datasets capturing transcriptome and methylome of fibroblasts and iNs. We defined the distinct cell states of *in vitro* cellular iN models and characterized epigenetic signatures of AD from age-retaining iNs. A comparative analysis between *in vitro* cellular models and *in vivo* primary brain tissues identified conserved and robust methylation signatures. A reliable set of machine learning model selected CpG sites showed a flawless accuracy of AD prediction across *in vitro* and *in vivo* cell types. In summary, we generated a comprehensive dataset dissecting the underlying molecular alterations involved in epigenetic regulation and 3D chromatin conformations of *in vivo* primary brain tissues and *in vitro* cellular iN models (Figure 1).

**Figure 1.**
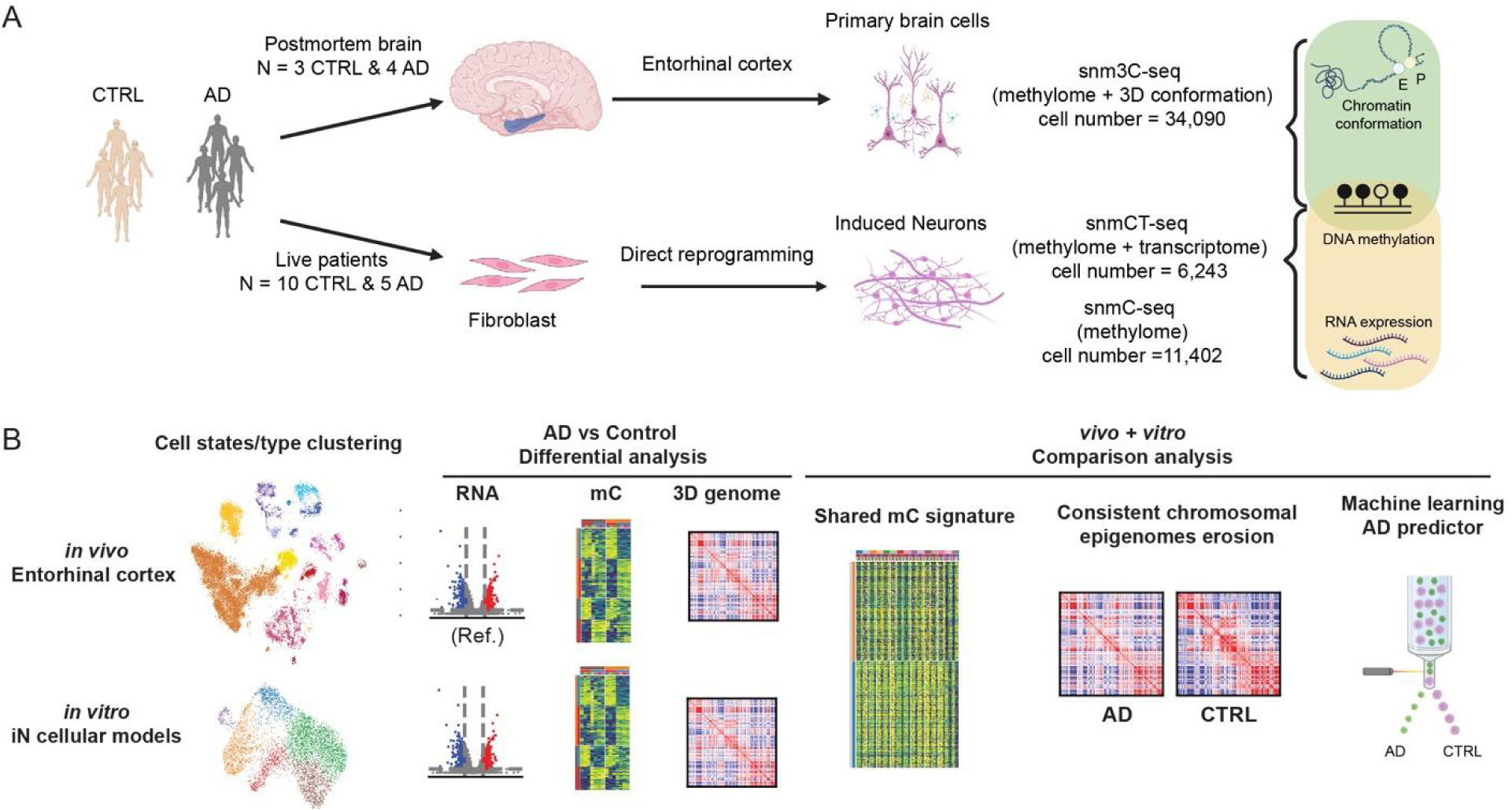
Overview of single-nucleus multi-omics to dissect epigenome erosion of LOAD in *in vivo* entorhinal cortex and *in vitro* iNs. (A) Schematic illustration of generating single-nucleus multi-omics datasets on postmortem entorhinal cortex and fibroblast-derived iNs. (B) Design to conduct AD versus CTRL differential analysis and *in vivo* versus *in vitro* comparison.

## RESULTS

### Identification of aDMRs in primary entorhinal cortex

The single nucleus RNA sequencing (snRNA-seq) and ATAC sequencing (snATAC-seq) of AD brain tissues have demonstrated that AD-specific transcriptome changes strongly depend on cellular identity.^12–15^ However, the alterations of DNA methylation and 3D chromatin architecture in LOAD brain cell types are still unclear. We applied our single nucleus multi-omics technologies, snm3C-seq^7^ capturing methylome and 3D chromatin conformation to 4 AD and 3 age-matched controls’ (CTRL) post-mortem human entorhinal cortex, a region critical in the development of AD^16,17^ (Supplementary Table 1). Collectively, 34,090 nuclei passed rigorous quality control, with 2.3 ± 0.7 million unique mapped reads and 4.3 ± 1.4 x e5 chromatin contacts detected per cell (Figure S1A and S1B). These provided reliable quantification of methylome and detection of active or repressive chromatin compartments, topologically associating domains (TADs), and chromatin loops in distinct cell types in the entorhinal cortex. After integration with human brain methylome datasets ^6^ based on mCH and mCG across 100-kb genomic bins, we annotated six cell classes, including excitatory neurons (Ex), inhibitory neurons (Inh), astrocytes (ASC), microglia (MGC), oligodendrocyte progenitor cells (OPC), and oligodendrocytes (ODC). They were further separated into 24 major cell types (Figure 2A and Methods). Excitatory neuron clusters were separated into cortical layers (L) (L2/3, L4, L5, L5/6, and L6) and projection types (IT = intratelencephalic, CT = corticothalamic, ET = extra telencephalic, and NP = near-projecting). Inhibitory neuron types expressing different neuropeptides could be separated, including Vip, Sst, and Pvalb, as well as rarer types like Lamp5_Lhx6 and Pvalb_ChC. A small fraction of CA2 neurons might be attributable to dissection contamination of the adjacent region between the hippocampus and entorhinal cortex. We observed no significant (p-value = 0.469, t-test) cell type proportion differences between 3 CTRL and 4 AD donors, with slightly increased ASC (p-value = 0.021) and decreased MGC proportions (p-value = 0.024) in AD individuals (Figure 1B and 1D). Consistent with findings in other vertebrate brain systems and young human donors^6^, 5mCs were also observed in abundance in non-CG (or CH, H= A, C, or T) contexts in aged human individuals, especially in neurons rather than glial cell types (Figure S1C). Global methylation levels across major cell types ranged from 74.2% to 81.9% for CG-methylation and 0.8% to 8.5% for CH-methylation without obvious differences across individuals. Consistent with the previous finding, gene activation negatively correlated with gene-body mCH levels at cell-type marker genes.^6,18,19^ For example, SATB2 is a marker gene for excitatory neurons and showed reduced mCH in excitatory neuron clusters (Figure 2C).

**Figure 2.**
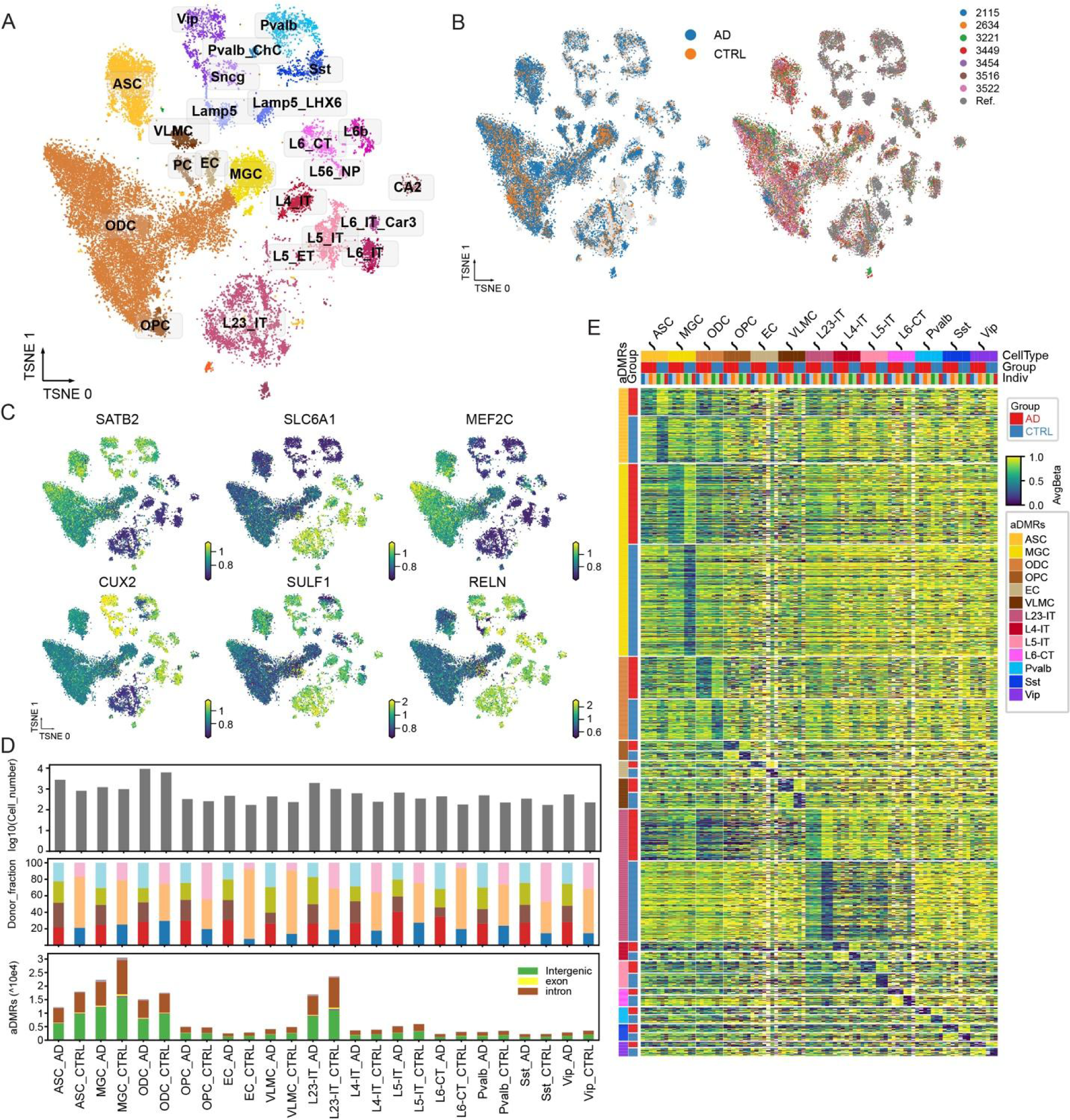
Epigenomic profiling of AD entorhinal cortex with snm3C-seq. (A) Iterative clustering and annotation of human brain nuclei. Clustering and visualization by t-distributed stochastic neighbor embedding (t-SNE) were based on mCG and mCH levels at 100kb bins. Major cell types were annotated and colored based on the integration analysis of methylome datasets from the Human Brain atlas published previously.^6^ (B) t-SNE visualization on the clustering colored by groups and individuals. (C) t-SNE visualization of gene body mCH levels for cell type marker genes, such as SATB2 (excitatory neurons), SLC6A1 (inhibitory neurons), MEF2C (all neurons), CUX2 (layer 2 and 3 neurons), SULF1 (oligodendrocytes), and RELN (astrocytes and oligodendrocytes). (D) The number of cell types, individual fractions and hypomethylated aDMRs. aDMRs bar plot was colored by genome features of the aDMRs located. For example, ASC_AD represents the aDMRs identified in astrocytes andhypomethylated in AD compared to CTRL. (E) Heatmap showing the average methylation level at aDMRs in pseudo bulk of major cell types across individuals. The columns were pseudo bulk samples grouped and colored by major cell types, groups (AD or CTRL) and individuals. The rows are hypomethylated aDMRs grouped and colored by finding cell types and hypomethylated groups.

To identify the AD-specific putative cis-regulatory elements in a brain cell type-specific manner, we grouped individuals into AD and CTRL and identified paired 209,972 aDMRs in the 13 major cell types (METHODS). MGC, ASC, ODC, and L2/3 IT neurons had the largest numbers of aDMRs located mostly at the intergenic and intronic regions (Figure 2D). The consistent methylation patterns of aDMRs across individuals showed robust AD/CTRL differences, most of which were cell-type specific (Figure 2E). Meanwhile, we also observed shared aDMRs across multiple cell types, like aDMRs located in gene GFPT2 introns, which were hypermethylated in AD L2/3 IT neurons and AD ODC (Figure S1D). GFPT2 (glutamine-fructose-6-phosphate transaminase 2) is involved in the glutamate metabolism pathway and controls the flux of glucose, which is becoming increasingly recognized as a hypometabolism phenotype in cancer and AD brain.^20,21^ In addition, we analyzed the TF motif enrichments (-log(p-value) > 10) in aDMRs across cell types (Figure S1E). For example, we identified the TF SPI1 enriched in aDMRs hypomethylated in CTRL microglia. The hypermethylated state of these aDMRs in AD microglia reflects the transcriptional repression in SPI1 target genes and is consistent with AD snATAC-seq analysis.^13^ Furthermore, we quantified the aDMRs distribution across the genome and identified 1,795 hotspots enriched for aDMRs across the autosomes (METHODS), 26% of these hotspots were shared between at least 2 major cell types (Figure 2A). The genome feature annotations of these hotspots showed a significant enrichment of CpG island and SINE-VNTR-Alu (SVA) class of retrotransposons (Figure 2B, Supplementary Table 2). Intriguingly, SVA as the evolutionarily young and hominid-specific retrotransposons and LINE1 are mobilized active in the human genome and are involved in human neurodegenerative diseases.^22,23^

### aDMRs interact with bivalent promoters of AD differential expression genes

To interrogate the multivalent interactions regulated by DNA methylation and 3D genome structures on transcriptional activity, we identified putative cis-regulatory elements (CREs) of differential expression genes (DEGs) identified in distinct cell types of snRNA-seq dataset^13^ by assigning the aDMRs to genes based on the loop interaction (METHODS). In total, 6,214 aDMRs/DEGs pairs, between 1,197 DEGs (across six major cell types) were assigned with 5,345 aDMRs in corresponding cell types (Supplementary Table 3). We found a significant enrichment of aDMRs at heterochromatin (Het) and zinc finger protein genes associated with chromatin states (Znf/Rpts)^24,25^ (Figure 3C and METHOD). These repressive states were usually marked by H3K9me3, associated with lamin-associated domains (LADs) and B compartments.^26–28^ However, the chromatin states on the promoters of DEGs linked with aDMRs across cell types show significant enrichment of repressed Polycomb states (ReprPC) and bivalent regulatory states (TssBiv and EnhBiv) (Figure 3D). The increased number of aDMRs linked to DEGs in excitatory neurons amplified the enrichment of these repressive states, like TssBiv, with depletion of active promoters (TssA1 and EnhA1) (Figure 3E). Bivalent promoters are usually marked both with active (H3K4me3) and repressive (H3K27me3) histone modifications.^29^ Approximately 15% of the promoters of total coding genes exhibit repressive bivalent states or are bound by the PRC complex in the human brain. This proportion surged to 50% for the promoters of DEGs linked with aDMRs in both excitatory and inhibitory neurons, as well as modest increments in glial cells (36% in ASC, 35% in ODC, and 24% in MGC) (Figure 3F and 3G). Examination of the combinatorial interactions between RNA expression of DEGs and average methylation alterations of linked aDMRs showed no clear correlation, suggesting a more complex relationship between epigenetic regulation and gene expression (Figure 3G). aDMRs linked DEGs in excitatory neurons are enriched in KEGG pathways related to glutamatergic synapse and axon guidance processes (Figure 3H). For example, one of the top genes linked with aDMRs in neurons, CSMD1 (CUB and Sushi Multiple Domains 1) was associated with 68 aDMRs in excitatory neurons and 10 aDMRs in inhibitory neurons and had bivalent promoter and upregulated RNA expression in AD samples (Figure 3G). A genetic variant of CSMD1 was previously found to be associated with schizophrenia and may be involved in the complement cascade system, including synaptic pruning and neuroinflammation.^30,31^ For certain DEGs in other cell types, like APOE in ASC, we identified 8 aDMRs associated with the gene, and 7 out of them were hypermethylated in AD samples, consistent with the downregulation of APOE RNA expression. An upregulated AD DEGs example in microglia and astrocytes, BCL6, had 12 aDMRs (MGC) and 5 aDMRs (ASC) linked via loop interactions. Eight of the twelve aDMRs interacting with the BCL6 gene in microglia were located in gene TPRG1 (Tumor Protein P63 Regulated 1), which is known to contain differentially methylated loci in epigenome-wide association studies (EWAS) of AD.^32,33^ One upregulated AD DEGs found in neurodegenerative disease and AD mouse models,^34^ HDAC4 (Histone Deacetylases 4) was upregulated specifically in ODC and interacted with 6 aDMRs spanning from 354 to 826 kbps downstream of the transcription start site (TSS), with three of them being hypomethylated in AD and three hypermethylated (Figure S2). In summary, our comprehensive datasets provide valuable resources to analyze the interplay between cis-regulatory elements and genes involved in AD pathogenesis, which enables us to characterize the associations of aDMRs with bivalent promoters and PRC repressive elements within DEGs.

**Figure 3.**
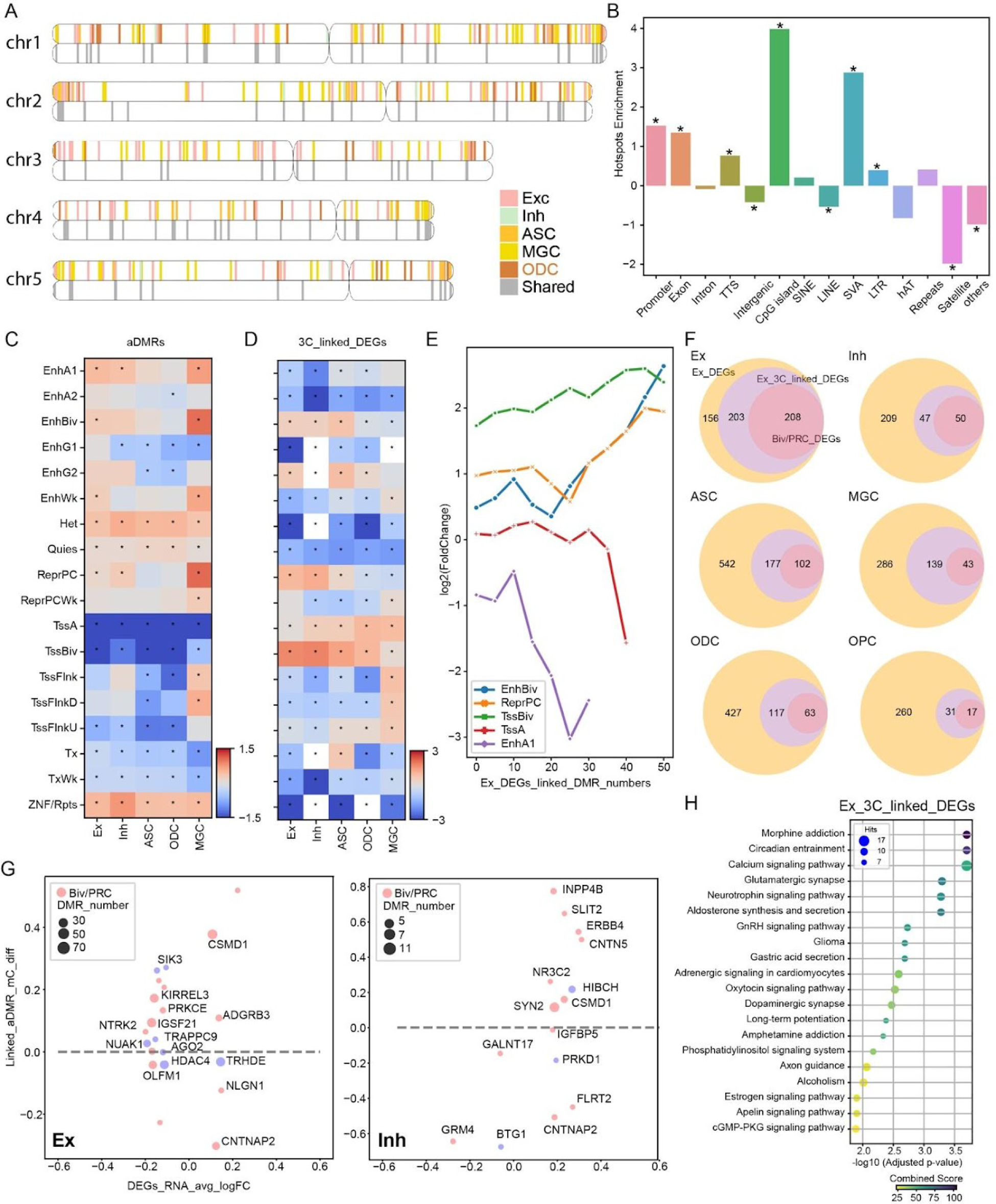
aDMR-enriched hotspots and interaction between aDMRs with repressive promoters of DEGs. (A) The distribution of 1,795 aDMR-enriched hotspots across the genomes, chr1-5 shown as examples. 5kb-bin hotspots were flanked to 1M for visualization and colored by the cell types where hotspots were identified from. For each chromosome, the top copy shows the cell type specific hotspots, the lower copy shows the shared hotspots found in at least two brain cell types. (B) Genome feature enrichments, Y-axis present log2 value of fold changes across the genome features (X-axis) between the aDMRs hotspots compared to the genome 5kb bin background. The star marked the p-value < 0.01 in fisher’s exact test. (C-D) Heatmap showing the log2 value of fold changes of ChromHMM states enrichment for the aDMRs (C) and the promoter of aDMRs linked DEGs in corresponding cell types (D). The star marked p-value < 0.01 in fisher’s exact test. (E) ChromHMM states enrichment analysis of the selected states (Repressive: EnhBiv, ReprPC, TssBiv; Active: TssA and EnhA1) on the promoter of aDMRs linked DEGs in excitatory neurons over the number of aDMRs linked. (F) Venn diagrams showing the overlapping of aDMRs linked DEGs, DEGs harboring promoters in repressive states (TssBiv and ReprPC), and DEGs from published snRNA-seq datasets.^13^ (G) Scatter plot of the average log2 RNA expression fold changes (AD/CTRL) of DEGs as X-axis and average methylation difference (AD-CTRL) of aDMRs linked to corresponding DEGs in excitatory and inhibitory neurons. The size and color represent the number of linked aDMRs and whether the promoter of the DEG is on repressive chromHMM states (TssBiv and ReprPC). (H) KEGG pathway enrichment analysis for aDMRs linked DEGs in excitatory neurons. X-axis shows the enrichment significance as −log (adjusted p-value), and Y-axis represents pathways. The size and color represent the number of related genes and combined score, respectively.

### Chromosomal epigenome erosion in AD brain cell types

Chromatin is organized into structures at different scales. The subchromosomal-level compartment brings together regions that are tens to hundreds of megabases (Mb) away, whereas TADs and chromatin loops are driven by interactions within several Mbs. Chromatin organization and related dysfunctional nuclear lamina (LMNA) in Hutchinson-Gliford Progeria have demonstrated the critical role of chromatin architecture in senescent cells, normal aging, and age-dependent disorders.^9,35–37^ The initial studies in neurodegeneration mouse models (Parkinson’s disease and AD) suggested abnormal dysfunctional histone modifiers such as SIRT and HDAC family.^38,39^ Further studies on heterochromatin protein 1α (HP1α), Polycomb group proteins, and ATP-dependent chromatin remodeler-like CHD5 indicated that the disrupted chromatin structures and organization contributed to aging and age-related neurodegenerative disorders.^40–42^ Chromatin accessibility assays in bulk (ATAC-seq) showed that the AD-associated cis-regulatory domains were enriched in A compartments.^43^ However, the 3D genome architecture and DNA loop contact maps in the LOAD brain, especially in distinct cell types, are still unknown.

We first analyzed the proportion of contacts detected at different genome distances within each single cell to examine the cell-type specificity of genome folding at different length scales. Within the same cell type, AD samples have significantly more longer-range interactions (20-50 Mb) and fewer shorter-range interactions (200 kb-2 Mb) compared to CTRL samples (Figures 4A to 4C, S3A and S3B). Given that megabase-level interactions are usually associated with compartment organization, we next investigated the relationship between enriched, longer-range chromatin contacts and chromatin compartments. We identified chromosome compartments in each cell type at 100 kb resolution and observed that enriched longer-range interactions in AD were predominantly inter-compartmental (Figure S3C). This finding differed from the enrichment of longer-range contacts seen in non-neural cells compared to neurons, where intra-compartment interactions dominated.^6^ Consistently, the compartment strength was weakened in AD, as quantified by the decrease in contact correlation between the intracompartment regions and the ratio between intracompartment and intercompartment interactions (Figures 4D to 4F and Figure S3D). We also identified domains at 25 kb resolution and loops at 10 kb resolution. The number of identified domains decreased in AD cells of all cell types (Figure 4G), whereas the number of loops significantly was reduced in AD samples only in ODC and MGC (Figure 4H). Generally, the insulation score at domain boundaries and the interaction strength at chromatin loops were weaker in AD (Figures S3D and S3E), consistent with the decreased shorter-range interactions. To associate the chromatin structure with gene expression, we first identified 43,620 differential loops (DL) between AD and CTRLs, with nearly all DL being lost in AD (99.8% of total DL). For example, TMEM59 is a transmembrane protein inhibiting APP transportation to the cell surface and downregulated in AD inhibitory neurons. Our data suggested that the chromatin loops surrounding the genes were also impaired during AD, and the DMRs associated with TMEM59 by chromatin loops also showed increased mCG (Figure 4G), suggesting the potential epigenetic dysfunction that led to the misregulation of its expression. Together, our data indicated that an erosion process of chromatin occurs in AD, which leads to the erroneous expression of bivalent repressive and PRC binding genes (Figure 4I and 4J).

**Figure 4:**
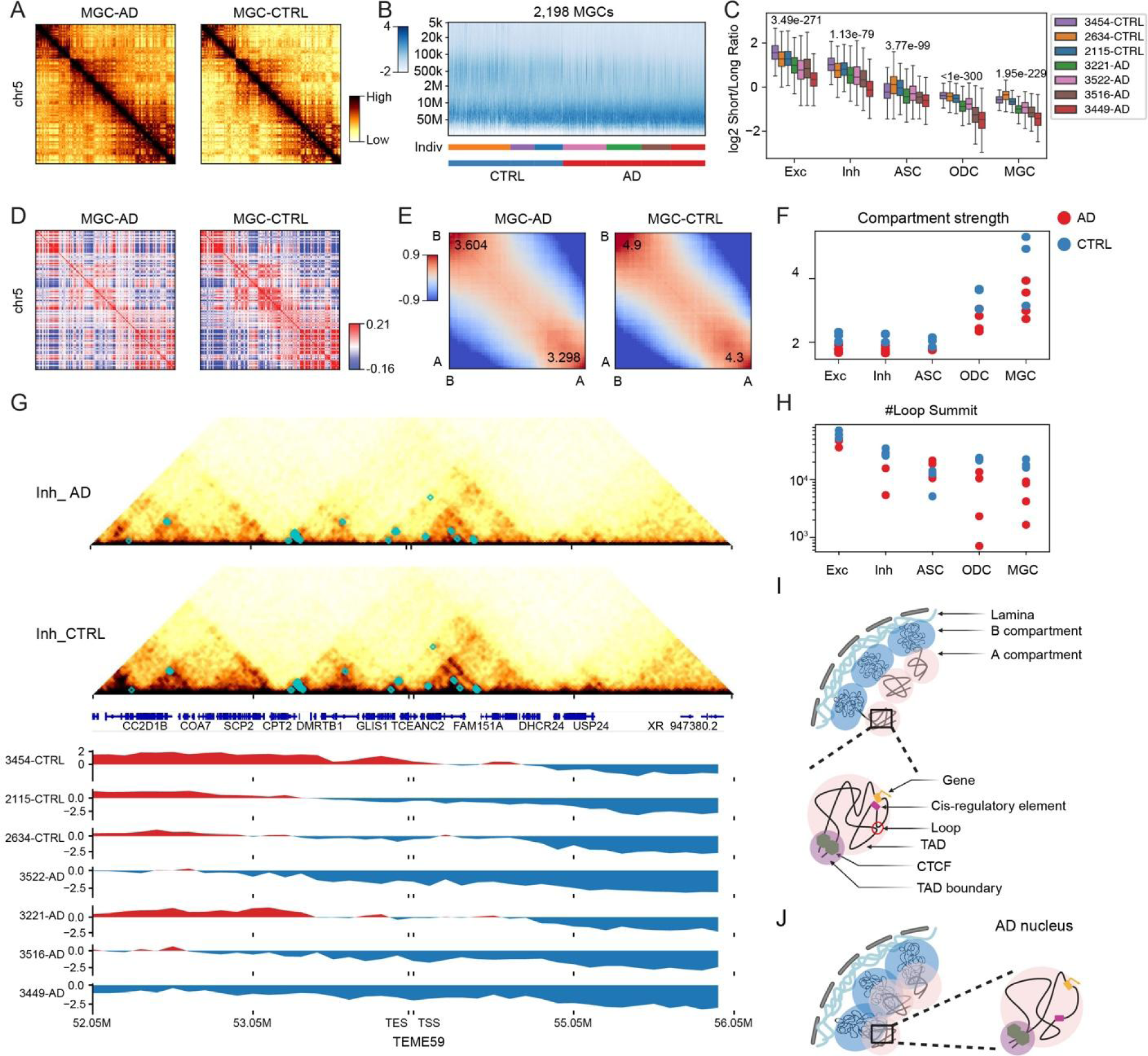
Chromosomal epigenome erosion in AD entorhinal cortex. (A) Chromatin contact map of AD and CTRL microglia at chromosome 5. (B) Frequency of contacts against genomic distance in each single cell of microglial type, Z-score normalized within each cell (column). The y-axis is binned at log2 scale. Each cell in the x-axis is grouped and colored by individuals and CTRL or AD. (C) The log2 short/long contacts ratio of major types across individuals. Centerline denotes the median, box limits denote the first and third quartiles, and whiskers denote 1.5 × the interquartile range. (D) Chromosome-wide Pearson’s correlation matrix of microglia cells from AD and CTRL. Chr5 as an example. Color bar ranges from −0.16 to 0.21. (E) Saddle plots (method) of microglial cells from AD and CTRL shown in (B), colored by contact frequency enrichment showing the interaction of A/B compartment in cis. (F) The compartment strength (AA+BB)/(AB+BA) across major cell types between AD (Red) and CTRL individuals (Blue). (G) Chromatin conformation around the gene TEME59 shows A to B compartment switching and decreased loop interactions in AD inhibitory neurons. Upper panel, chromatin contact map of AD and CTRL inhibitory neurons with differential loops between AD and CTRL marked in cyan box. Lower panel, A(Red) and B(Blue) compartments track across individuals (H) The total loop interaction numbers of major types across individuals. (I-J) Illustration of 3D chromatin conformation in normal brain cell types (I) and AD (J).

### snmCT-seq characterization of distinct cell states in human neurons directly converted from AD fibroblasts

Modeling age-dependent neurodegenerative diseases is by far one of the biggest challenges for researchers seeking to find cellular model systems or animals that can recapitulate temporal dynamics of up to years in duration. Reprogramming patient tissues to induced pluripotent stem cells (iPSCs) is a powerful approach for genetic-based disease modeling; iNs can be generated through differentiation from reprogrammed iPSCs or by direct conversion from patient somatic fibroblasts.^44,45^ However, the transition through the stem cell intermediate phase leads to a youthful rejuvenation of the epigenome (METHOD and Figure S4), gene expression, long-lived proteins, mitochondria function, and telomere length in the resulting neuron.^46–50^ The directly reprogrammed neurons from fibroblasts retain biological aspects of age ^49,51^ and disease.^52–54^ Single-cell RNA-seq studies in mouse iNs directly converted from fibroblasts have suggested there is cell state diversity during transdifferentiation.^55^ Genome-wide methylation and open chromatin dynamics revealed epigenome and chromatin reconfiguration during mouse direct reprogramming.^56,57^ However, the heterogeneity and epigenome dynamics of iN conversion of human fibroblasts, especially from aged patients and AD donors, have yet to be investigated.

To characterize the cell state transitions along the human fibroblast-to-iN conversion process and evaluate whether direct trans-differentiation iN models can mimic aging and AD signatures in primary brain tissues, we profiled iNs from 6 LOAD and 4 age-matched, cognitively normal control individuals, generating a snmCT-seq dataset of 6,242 cells as well as a snmC-seq dataset of 11,402 cells (Supplementary Table 1). The cells were not sorted with PSA-NCAM because we used snmCT-seq and can identify cells during analysis. The data quality is comparable to our previous publication ^8^ (Figure S5A; S5B). The methylome modality in the snmCT-seq dataset covered 3.47%± 1.7% (mean±s.d.) of the genome, whereas the transcriptome detected 4,273±1726 genes from each single nucleus. Based on transcriptome clustering, we identified 6 distinct cell states during the iN-induction process: fibroblast, mesenchymal to epithelial transition (MET), intermediate neuronal progenitor (INP), iN, Misc1 and Misc2 (Figure 5A and 5B). The downregulated expression of fibroblast marker genes (VIM, FN1, and CD44) and upregulated expression of neuron marker genes (MAP2, NCAM1, and CAMK4) from fibroblast to iN state confirmed the successful iN conversion in both AD and CTRL individuals (Figure 5C, 5D, Supplementary Table 4). The diversity of conversion efficiency across individuals can be quantified by the proportion of iN states after 3 weeks of the iN-induction process, varying from 22%± 3% in CTRL and 23%± 11% in AD (Figure S5E and S5F). Misc1 and Misc2 clusters have mixed marker genes and are labeled as uncharacterized cell states. Misc1 may be due to transition failure during the transdifferentiation since several genes within heterochromatin are active, like CSMD gene families. In contrast, cell cycle-related genes like CENP family genes and DNA replication genes are highly expressed in the Misc2 cluster, suggesting it may represent cells in oncogenesis or senescence (Supplementary Table 4).

**Figure 5.**
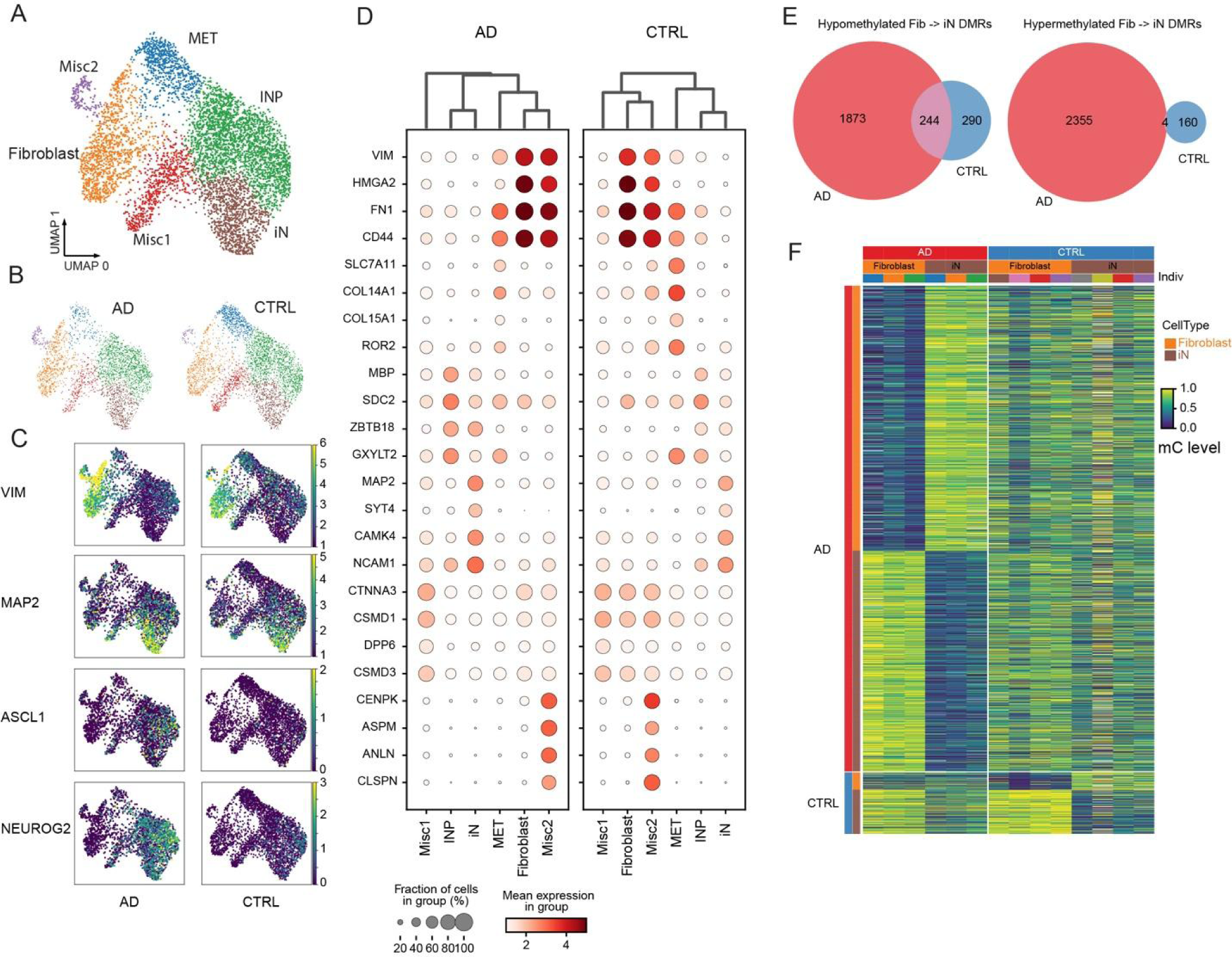
Single nucleus methylome and transcriptome sequencing (snmCT-seq) characterizing distinct cell states in direct converted neurons from fibroblasts. (A) Two-dimensional UMAP visualization of snmCT-seq data for fibroblasts and direct-converted iNs. Each dot represents a nucleus colored by annotated cell states. (B) Nucleus clustering colored by cell states in AD and CTRL. (C) UMAP of nucleus colored by logarithmized CPM of marker gene VIM (fibroblasts), MAP2 (iNs) and two proneuronal factors ASCL1 and NGN2. (D) The top RNA marker genes in each specific cell state in AD and CTRL. (E) Venn diagrams of shared Fibroblast −> iN conversional DMRs between AD and CTRL. Hypomethylated Fibroblast->iN DMRs presenting the DMRs loss of methylation from fibroblasts to iNs and hypermethylated means gain of methylation during iN induction. (F) Heatmap showing the average methylation fraction at conversional DMRs in pseudo bulk of fibroblasts and iN cell states across individuals. The columns are pseudo bulk samples grouped and colored by group (AD or CTRL), cell states (fibroblasts and iNs) and individuals. The rows are hypomethylated conversional DMRs grouped and colored by group (AD or CTRL) and cell states (fibroblasts and iNs).

To examine methylome reconfiguration during iN induction, we compared fibroblasts within populations from the same individuals, i.e., CTRL and AD cells were compared separately. In total, 4,476 (AD) and 698 (CTRL) fibroblast −> iN conversion-related DMRs (cDMRs) were identified and were consistent across individuals within the AD/CTRL groups (Figure 6E). We observed 244 cDMRs that were shared between AD and CTRL groups, and almost all of them had methylation levels decreased from fibroblast to iN, suggesting that the overexpression of ASCL1/NGN2 neuronal factors initiated a shared demethylation program at downstream targets regardless of disease status. Conversely, only 4 cDMRs that gained methylation during iN induction were shared between AD and CTRL. Overall, cDMRs were in intergenic (51.67%± 11.08%) and intronic (41.19%± 11.38%) genome features (Figure S5G). The motif enrichment analysis of cDMRs by HOMER^58^ revealed a shared demethylation pattern surrounding the binding sites of Zinc fingers (ZFs) and basic Helix-Loop-Helix (bHLH) TFs; there was no significant (p-value < 0.01) motif enrichment at hypermethylated regions along the iN conversion trajectory (Figure S5H). Intriguingly, neuronal differentiation genes like TCF4 coupling with induction factors ASCL1 and NGN2 showed increased RNA expression and their binding motifs are enriched at the cDMRs demethylated during the iN conversion process (Figure S5I). Additionally, we observed a higher expression level of ASCL1 and NGN2 and more cDMRs found during AD iN induction, suggesting that the epigenetic landscape of AD fibroblasts is more permissible for iN transdifferentiation. In summary, both AD and CTRL fibroblast-derived iNs can be generated successfully with a shared ASCL1/NGN2-initiated demethylation process, and 6 cell states can be observed both in AD and CTRL during transdifferentiation.

**Figure 6:**
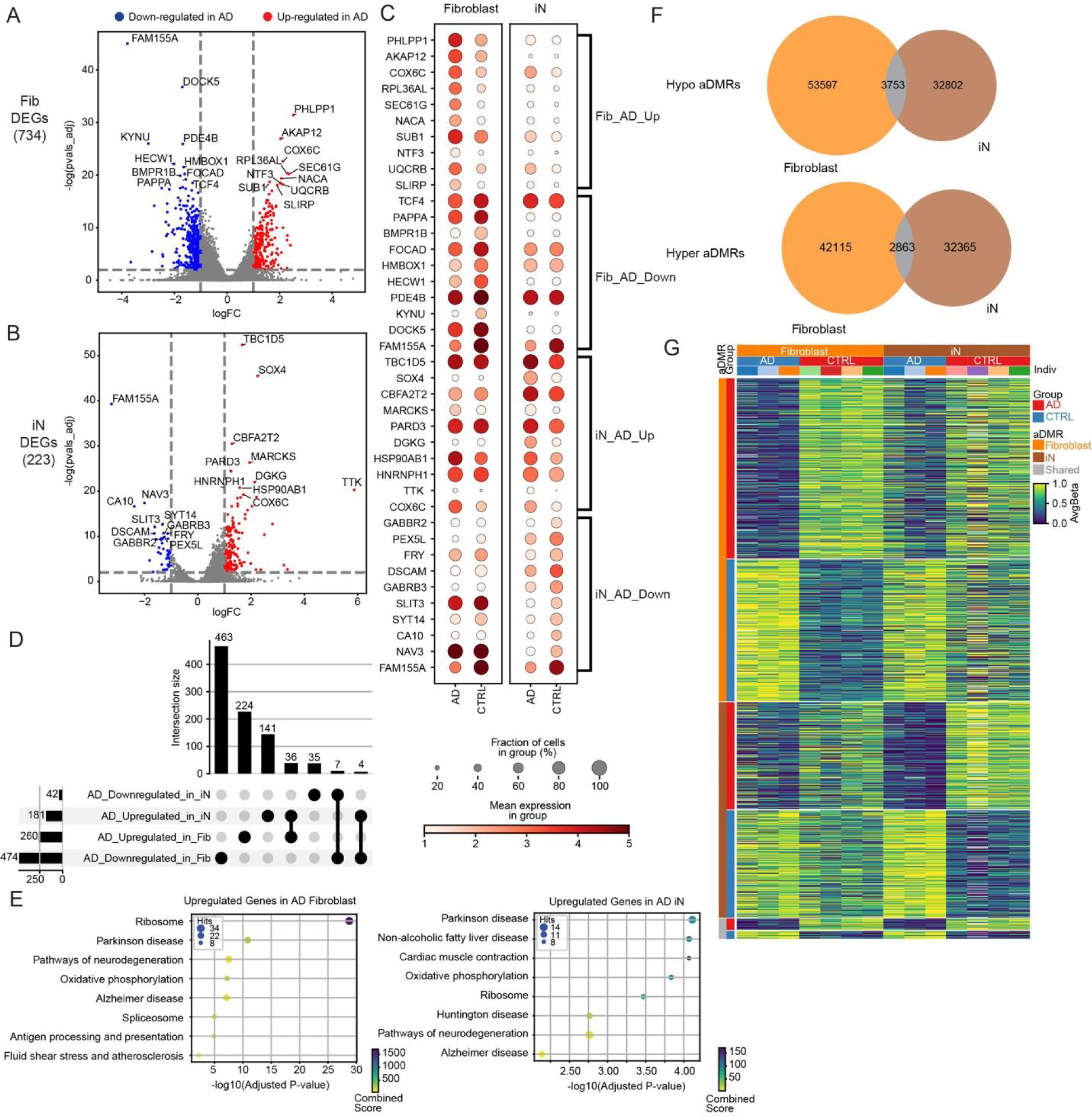
Joint analysis of DNA methylome and RNA identifies DEGs and aDMRs in human fibroblasts and fibroblast-induced neurons. (A-B) DEGs between AD versus CTRL in fibroblast (A) and iN (B) states, colored by up- (Red) or down-regulated (Blue) DEGs in AD samples. (C) The top up- or down-regulated DEGs in fibroblasts and iNs. (D) UpSet plots summarized up- or down-regulated DEGs in fibroblasts and iNs. The bottom left horizontal bar graph shows the total number of DEGs. The top bar graph presents unique or overlapping DEGs. (E) KEGG pathway enrichment analysis for up-regulated DEGs in fibroblasts and iNs. X-axis shows the enrichment significance as −log(adjusted p-value), and Y-axis represents pathways. The size and color represent the number of related genes and combined score, respectively. (F) Venn diagrams showing the overlapping of aDMRs in both fibroblasts and iNs. For example, hypomethylated aDMRs present the aDMRs hypomethylated in AD whereas hyper aDMRs means hypermethylated in AD samples. (G) Heatmap showing the average methylation level at aDMRs in pseudo bulk of fibroblast and iN cell states across individuals. The columns are pseudo bulk samples grouped and colored by cell states (fibroblasts and iNs), groups (AD or CTRL) and individuals. The rows are hypomethylated aDMRs grouped and colored by cell states (fibroblasts and iNs) and group (AD or CTRL).

### AD-specific methylation and transcriptome signatures in fibroblast-induced neurons

We compared AD versus CTRL groups within fibroblast and iN states to identify AD-specific transcriptomic and methylation signatures. To this end, we identified 734 DEGs (p-value < 0.01, log2foldchange >1) between AD and CTRL in fibroblasts and 223 in iNs. Thirty-six (7) DEGs were upregulated (downregulated) in AD in both fibroblast and iN states (shared DEGs; Figure 6A-D, Supplementary Table 5). We observed that 9 of the 36 shared DEGs upregulated in AD belonged to the ATP synthesis pathway in mitochondria, for example, UQCRB and COX6C. One of 7 AD-downregulated shared DEGs, FAM155A, was also downregulated in AD in endothelial cells.^59^ Strikingly, GABA receptors (e.g., GABBR2 and GABRB3) together with synaptic transmission-related genes, KCND2 and SYT14, were only observed to be downregulated in AD in iNs rather than fibroblast (Figure 6C and Supplementary Table 5). KEGG pathway enrichment analysis revealed that the pathways related to neurodegeneration were enriched in AD-upregulated genes both in fibroblasts and iNs. In contrast, genes involved in synaptic transmission and neuron development were significantly downregulated only in AD iNs (Figure 6E and Supplementary Table 5).

Regarding the methylome, we identified 160,879 aDMRs, most distinct in either fibroblast or iN. We detected 3,753 (2,863) aDMRs hypomethylated (hypermethylated) in AD shared between fibroblast and iN states (Figure 6F). These aDMR patterns depicted distinct DNA methylation signatures between AD and CTRL for fibroblast and iN identities (Figure 6G). To gain insights into the potential impact of DNA methylation on the binding of TFs, we conducted motif analysis on these aDMRs, revealing 4 top TF families (ETS, ZFs, bHLH, and bZIP) with a significant score (-log(p-value) > 15). Most ETS TFs were assigned to hypomethylated DMRs in CTRL fibroblasts and iNs. In contrast, bHLH TF enrichments were specific to fibroblast cell states, and bZIP TF enrichments were specific to aDMRs hypomethylated in CTRL fibroblasts (Figure S6A). By examining the RNA expression of these TFs from the same cells, we narrowed down the putative TF candidates within each TF family and dissected the interactions between expressing TF and methylation states of their binding motifs (Figure S6B). For instance, the AD upregulated TFs, BHLHE40 and its close homolog BHLHE41, have specific enrichments at hypomethylated aDMRs in AD fibroblast, consistent with their binding preference of unmethylated CpG sequences. ^60^ Both TFs are crucial in the regulation of cholesterol clearance and lysosomal processing in microglia and may be associated with AD pathogenesis.^61,62^

Furthermore, we integrated information on DEGs and DMRs to identify putative CREs in fibroblasts and iNs. DMRs were associated with genes by GREAT algorithms.^63^ In total, 10,070 aDMRs were paired with 659 DEGs in fibroblasts, and 2,963 aDMRs were identified associated with 197 DEGs in iNs. The RNA expression changes of these DEGs and methylation alteration of the associated aDMRs revealed an orchestrated gene regulation program between AD and CTRL *in vitro* cellular models (Figure S6C and S6D). For example, the downregulated shared DEG, FAM155A, was associated with 130 aDMRs in fibroblast and 83 aDMRs in iN that were hypomethylated in AD on average. In contrast, 63(35) aDMRs in fibroblast (iN) assigned to the up-regulated PAX3 gene were hypermethylated in AD samples. Concordantly, EWAS of AD has reported PAX3 to be a hotspot, showing hypermethylation in AD hippocampus.^64^ Systematic examination of RNA fold changes of DEGs and methylation difference of associated aDMRs revealed a positive correlation and the correlation coefficient increased with the number of associated aDMRs (Figure S6E). This positive correlation between gene expression and mCG had been observed previously in cultured cell lines as a depletion of methylation in repressive genes located in partially methylated domains (PMDs), resulting in the reactivation of gene expression.^65^

### Shared aDMRs between the *in vitro* fibroblast/iN model and isogenic entorhinal cortex cell types

There has been no systematic comparison on genome-wide epigenome between *in vitro* iN modeling and *in vivo* primary brain tissues from isogenic patients. We had profiled both the entorhinal cortical tissue and cultured fibroblasts and iNs of 2 AD and 1 CTRL donors. Overall, there is a small fraction (3,733) of aDMRs overlapping between *in vitro* fibroblast/iNs and *in vivo* entorhinal cortex; 2,274 of them have the same direction of methylation changes and are consistent across individuals and *vitro*/*vivo* cell types (Figure 7A and 7B). To validate the reliability of these aDMRs and investigate individual differences, we did a random sampling of aDMRs from *in vitro* or *in vivo* aDMR pools. We compared them to the 2,274 aDMRs between the 2 AD patients and 1 CTRL donor. Distinguishable methylation patterns were evident in the shared aDMRs, whereas those identified from either *in vitro* or *in vivo* methods lacked consistency (Figure 7C). The motif enrichment analysis on these shared DMRs revealed putative TFs candidates (Figure S7A), like pluripotency associated TF, Pou5F1 (OCT4), as well as early growth response-1 (EGR1) which have already been found to play crucial roles in early stages of AD,^66,67^ are specifically enriched in the hypomethylated aDMRs in AD samples.

**Figure 7:**
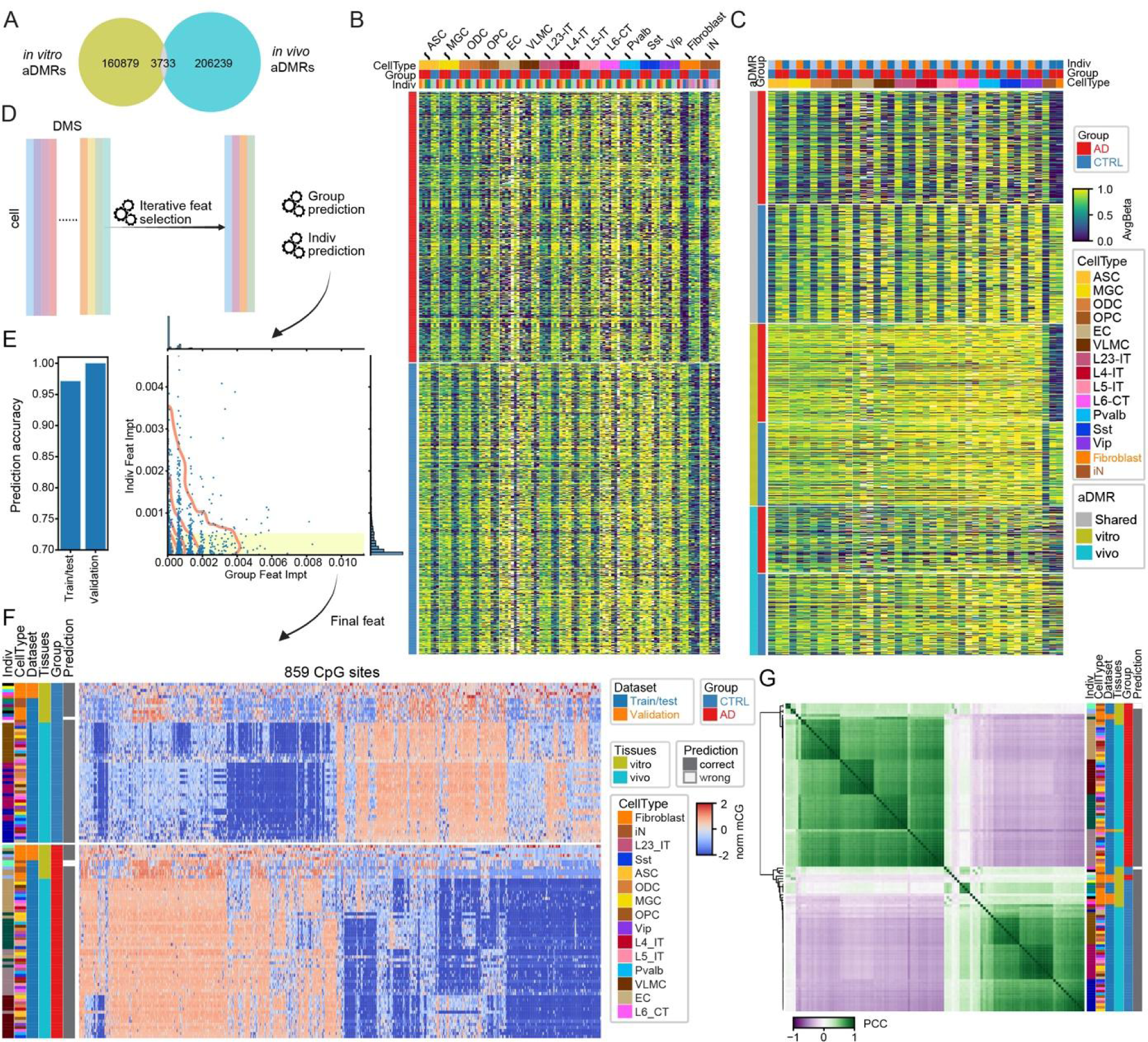
A methylome-based predictive model captures AD-specific DNA methylation signatures in *in vitro* Fibroblast/iNs and *in vivo* entorhinal cortex brain cell types. (A) Venn diagrams of shared aDMRs between *in vitro* iN cellular models and *in vivo* primary entorhinal cortex. (B) Heatmap showing the average methylation level at shared aDMRs (1,095 hypomethylated and 1,179 hypermethylated in AD) in pseudo bulk of major cell types including *in vivo* brain cell types and *in vitro* fibroblast/iNs across individuals. (C) Heatmap showing the average methylation level in isogenic individuals have both *in vitro* and *in vivo* cell types at shared aDMRs, and random selected aDMRs identified only from *in vitro* or *in vivo* systems (aDMRs number: 2,274 shared, 2,207 random sampling from 160,879 *in vitro* aDMRs, 1,798 random sampling from 206,239 *in vivo* aDMRs) (D) Design of methylome predictors across *in vitro* fibroblast/iN and entorhinal cortex tissues. (E) The bar plot shows the accuracy of AD predictors used in validation for cross-individuals. (F) Heatmap presents the normalized mC level at the selected features of AD predictors across the individuals, groups and cell types. The prediction results were marked as correct in gray and wrong in white. (G) The Pearson coefficient correlation shows the similarity of the pseudo bulk methylation pattern at selected features across individuals, groups and cell types.

### A reliable set of differentially methylated sites for AD prediction

Both cell type variations and individual differences influence the methylation fractions of differentially methylated sites (DMSs) between AD and CTRL groups. To identify DMSs consistent across cell types and resilient to individual variability, we devised a machine learning (ML) method (Figure 7D). A meticulous iterative feature selection was employed to hone in on candidate CpG sites (see Methods), resulting in a prediction accuracy of 97.1% to distinguish AD v.s. CTRL donors (Figure 7E). We further filtered out sites influenced by individual effects (METHODS) to enhance the prediction robustness, ultimately pinpointing 859 CpG sites. Although samples exhibited variances between *in vivo* and *in vitro* conditions, the methylation patterns of these sites consistently differentiated AD from the CTRL group, remaining stable across diverse cell types and individuals (Figure 7F, 7G, and Supplementary Table 6). Remarkably, when trained on *in vivo* snm3C datasets combined with *in vitro* snmCT datasets and applied to a separate validation dataset (Figure S7B and Supplementary Table 1) using different technologies (snmC-seq, only methylome modality) comprising shared individuals and three unseen donors, our prediction model, grounded on these sites, achieved a flawless accuracy of 100%. Given the robustness of these sites and the predictive model across cell types and donors, they promise valuable insights for subsequent research. The ML-selected CpG sites also have biological importance. For example, 8 ML-selected CpG sites around the BCL6 gene overlapped with aDMRs-enriched TPRG1 and P3H2 genes (Figure S7C). BCL6 is one of the DEGs in microglia and astrocytes, whereas the TPRG1 locus comprises aDMRs that could work as CREs of the BCL6 gene via loop interactions.

## Discussion

The intricacy of age-related dependencies in human brain neurodegeneration is challenging to recapitulate in cell models, significantly constraining our investigation of the molecular mechanisms underlying LOAD pathogenesis. Direct reprogrammed neurons were confirmed to retain aging and AD transcriptomic signatures, providing a novel approach to dissect the biological events occurring in AD brains. For *in vitro* cellular iN models, we characterized six distinct cell states along the iN-induction process. We found that AD fibroblast nuclei had a more permissive epigenetic state for ectopic expression of induction factors. We also identified the DEGs and DMRs between AD and CTRL in a cell state-specific manner, which are potential molecular biomarkers in the iN model and can be further explored in the functional studies of AD pathogenesis.

In addition, we generated the first single-cell multi-omics datasets capturing methylome and 3D chromatin conformation on the entorhinal cortex from LOAD patients. Moreover, we compared brain cell types with isogenic *in vitro* cultured fibroblasts and iNs derived from the same brain donors. Based on the comparison between the *in vitro* cellular model and *in vivo* primary brain tissues, we identified robust aDMR candidates that demonstrated consistency across cell types and individuals. Utilizing ML algorithms using these datasets, we developed a minimum and reliable prediction model to conduct LOAD diagnosis on *in vitro* cellular fibroblast/iN models and primary brain tissues. Moreover, we found global chromosomal epigenome erosion in brain cell types from LOAD donors, consisting of disrupted active/repressive chromatin compartments, weakened chromatin domain boundaries, and decreased short DNA loop interactions. Furthermore, by integrating our data with published snRNA-seq datasets on LOAD patients, we identified the potential CREs interacting with the DEGs involved in the AD process at a cell type-specific level. These findings suggest that an age-dependent dysfunctional genome architecture in brain cell types plays a fundamental role in neurodegeneration. In addition to the current molecular brain cell atlas efforts in the BICCN consortium,^6^ our datasets provide a comprehensive landscape and pilot study to phenotype the epigenome of the aging brain and AD cognitive disorders.

### Limitations

Like other plate-based, genome-wide single-cell multi-omics assays, the relatively high genome coverage and sequencing cost limit the number of cells and individuals profiled. However, with the new generation short reads sequencer continually reducing sequencing costs, our technologies and findings will become feasible to be validated in a larger group. Nevertheless, recent studies on the methylation clock and EWAS suggest that the methylome and epigenetic regulations play a key role in aging and age-dependent neurodegeneration disorders like LOAD. We observed that part of the aDMRs overlapped with these microarray-based methylated loci found in large cohorts; it will be necessary to see how the robustness and consistency of our identified aDMRs and ML-selected loci working in a large population in either brain cell types or fibroblast/iN cellular models. Although we revealed the phenotype of chromosomal epigenome erosion using sequencing-based snm3C-seq technology, the imaging approach may be another way to confirm the global chromatin alterations in AD brain cell types, at least in some locus-specific manner. Nevertheless, our findings have generated insights and provided pilot candidates to study the interplay between and among methylome, transcriptome, and 3D chromosome for AD pathogenesis in both *in vitro* cellular models and *in vivo* primary brain cell types, suggesting our datasets’ unique and constructive value for multiple fields.

## Lead contact

Further information and requests for resources and reagents should be directed to and will be fulfilled by the Lead Contacts Fred H. Gage and Joseph R. Ecker.

## Data and code availability

Our raw data is under uploading to GEO. The pseudo bulk methylation, RNA bams and aDMRs tracks can be accessed in a browser as well.

## METHODS

### Ethics statement and IRB clinical information of AHA-Allen cohort

Based on the clinical criteria published by the Consortium to Establish a Registry for Alzheimer’s Disease (CERAD), National Institutes of Health (NIH) standards, and Braak staging, subjects in AHA-Allen cohorts were recruited by Shiley-Marcos UCSD Alzheimer’s Disease Center. Dermal human fibroblasts and postmortem entorhinal cortex were collected with informed consent and strict adherence to legal and ethical guidelines from patients of the Shiley-Marcos UCSD Alzheimer’s Disease Center.

### Human iPSC lines and generation of iNs

Human iPSCs were obtained from the Salk Stem Cell Core. Fibroblasts were reprogrammed via CytoTune™-iPS 2.0 Sendai Reprogramming Kit (ThermoFisher Cat # A165167) per manufacturer recommendations. All iPSCs were karyotypically validated via g-banded karyotyping (WiCell) and are regularly screened for mycoplasm via MycoAlert™ PLUS Mycoplasma Detection Kit (Lonza Cat # 75860-362). Two major approaches to generating neurons in a dish are based on overexpression of proneural factors combined with chemicals from iPSCs differentiation or directly converted from fibroblasts. Age-dependent transcriptional signatures are more likely to be retained in directly converted neurons from fibroblasts rather than in differentiated iPSCs.^51^ To assess different differentiation strategies to generate iNs and characterize their epigenome modality, we conducted single nucleus methylome sequencing (snmC-seq) to profile the methylome of iN cells differentiated from either human pluripotent stem cells or human fibroblast cells via the overexpression of proneuronal factor NGN2 approach. Using young (1 yr. old, male) and aged (76 yrs. old, male) fibroblasts as cell resources, we found that iNs generated from fibroblasts retained aging methylation features and individual differentially methylated region (DMR) signatures, whereas the iPSC-iN method did not. DMRs were erased during iPSC reprogramming and reconfigured during NPC differentiation, and iPSC-iN cells from young and aged samples became indistinguishable (Figure S4). Direct conversion of iNs was performed via doxycycline-inducible NGN2-P2A-ASCL1 (N2A) as previously described.^51^ Briefly, stable N2A fibroblast lines were generated with lentivirus. Fibroblasts were maintained in dense cultures and passaged three times under puromycin selection before induction. Upon confluence, media was replaced with neural conversion media containing doxycycline for 21 days. For single-nuclei experiments, iNs were washed with PBS, incubated for 20 minutes at 37°C with TrypLE (Gibco cat #12604039), diluted in PBS up to 15 mL, pelleted at 100 x g for 5 minutes, aspirated, and snap frozen.

### Nuclei purification from *in vivo* fibroblasts and iNs cells for snmCT-seq

Cultured fibroblast and induced iN cells in the dish were dissociated in TrpLE medium. Cells were counted and aliquoted at 1 million per experimental sample and then pelleted by centrifugation at 100 x g for 5 min. The supernatant medium was aspirated, and cell pellets were resuspended in 600 μl NIBT [250 mM Sucrose, 10 mM Tris-Cl pH = 8, 25 mM KCl, 5mM MgCl2, 0.1% Triton X-100, 1 mM DTT, 1:100 Proteinase inhibitor (Sigma-Aldrich P8340), 1:1000 SUPERaseIn RNase Inhibitor (ThermoFisher Scientific AM2694), and 1:1000 RNaseOUT RNase Inhibitor (ThermoFisher Scientific 10777019). After gently pipetting up and down 40 times, the lysate was mixed with 400 ml of 50% Iodixanol (Sigma-Aldrich D1556) and loaded on top of a 500 ml 25% Iodixanol cushion. Nuclei were pelleted by centrifugation at 10,000 x g at 4°C for 20 minutes using a swing rotor. The pellet was resuspended in 2 mL of DPBS supplemented with 1:1000 SUPERaseIn RNase Inhibitor and 1:1000 RNaseOUT RNase Inhibitor. Hoechst 33342 was added to the sample to a final concentration of 1.25 nM and incubated on ice for 5 minutes for nuclei staining. Nuclei were pelleted by 1,000 x g at 4°C for 10 minutes and resuspended in 1 mL of DPBS supplemented with RNase inhibitors.

### snmCT-seq library preparation

The optimized snmCT-seq library preparation is based on the snmCAT-seq published previously^8^. A detailed bench protocol can be found at https://www.protocols.io/view/snmcat-v2-x54v9jby1g3e/v2. In general, the purified nuclei were sorted into a 384-well plate (ThermoFisher 4483285) containing 1 μl mCT reverse transcription reaction per well. The mCT reverse transcription reaction contained 1X Superscript II First-Strand Buffer, 5 mM DTT, 0.1% Triton X-100, 2.5 mM MgCl2, 30 mM NaCl, 500 mM each of 50-methyl-dCTP (NEB N0356S), dATP, dTTP and dGTP, 1.2 mM dT30VN_5 oligo-dT primer, 2.4 mM TSO_4 template switching oligo, 2 mM N6_3 random primer, 1U RNaseOUT RNase inhibitor, 0.5 U SUPERaseIn RNase inhibitor, and 10 U Superscript II Reverse Transcriptase (ThermoFisher 18064-071). The plates were placed in a thermocycler and incubated using the following program: 25 °C for 5 minutes, 42 °C for 90 minutes, 10 cycles of 50 °C for 2 minutes and 42 °C for 2 minutes, 85 °C 5 minutes followed by 4 °C. Three μl of cDNA amplification mix was added into each snmCT-seq reverse transcription reaction. Each cDNA amplification reaction contained 1X KAPA 2G Buffer A, 600 nM ISPCR23_3 PCR primer, and 0.08U KAPA2G Robust HotStart DNA Polymerase (5 U/mL, Roche KK5517). PCR reactions were performed using a thermocycler with the following conditions: 95 °C 3 minutes −> [95 °C 15 seconds −> 60 °C 30 seconds −> 72 °C 2 minutes] −> 72 °C 5 minutes −> 4 °C. The cycling steps were repeated for 12 cycles. One μl uracil cleavage mix was added into cDNA amplification reaction. Each 1 μl uracil cleavage mix contained 0.5 μl Uracil DNA Glycosylase (Enzymatics G5010) and 0.5 μl Elution Buffer (QIAGEN 19086). Unincorporated DNA oligos were digested at 37 °C for 30 minutes using a thermocycler. After addition of 25 μl of conversion reagent (Zymo Research) was added to each well of a 384-well plate, the following bisulfite conversion and library preparation was based on snmC-seq2 (described previously^68^) and on an updated version snmC-seq3 used in BICCN.^69^

### Nuclei purification from human postmortem tissues for snm3C-seq

Brain blocks were ground in liquid nitrogen with cold mortar and pestle and then aliquoted and stored at −80 °C. Approximately 100 mg of ground tissue was resuspended in 3 mL NIBT as above. The lysate was transferred to a pre-chilled 7 mL Dounce homogenizer (Sigma-Aldrich D9063) and Dounced using loose and tight pestles for 40 times each. The lysate was then mixed with 2 mL of 50% Iodixanol (Sigma-Aldrich D1556) to generate a nuclei suspension with 20% Iodixanol. One ml of the nuclei suspension was gently transferred on top of a 500 ml 25% Iodixanol cushion in each of the 5 freshly prepared 2-ml microcentrifuge tubes. Nuclei were pelleted by centrifugation at 10,000 x g at 4 °C for 20 minutes using a swing rotor. The pellet was resuspended in 1ml of DPBS supplemented with 1:1000 SUPERaseIn RNase Inhibitor and 1:1000 RNaseOUT RNase Inhibitor. A 10-μl aliquot of the suspension was taken for nuclei counting using a Biorad TC20 Automated Cell Counter. One million nuclei aliquots were pelleted by 1,000 x g at 4 °C for 10 minutes and resuspended in 800 μl of ice-cold DPBS.

### snm3C-seq library preparation

The purified nuclei for snm3C-seq were cross-linked with additional digestion and ligation to capture *in situ* long-range DNA interaction following a modified protocol of Arima-3C kit (Arima Genomics). A detailed bench protocol can be found in the BICCN atlas paper.^69^

### Automation and Illumina sequencing

The prepared nuclei from either snmCT-seq or snm3C-seq were sorted into a 384-well plate by Influx (BD) on a one-drop single mode. Then the automation handling of plates and library preparation for both snmCT-seq and snm3C-seq libraries followed the same bisulfite conversion-based methylation sequencing pipelines described previously ^18,68^ and an updated version snmC-seq3 used in BICCN.^69^ To facilitate large-scale profiling, Beckman Biomek i7 instrument was used and running scripts were shared.^69^ The snm3C-seq, snmCT-seq and snmC-seq libraries were sequenced on an illumina NovaSeq 6000 instrument using one S4 flow cell per 16 384-well plates on 150-bp paired-end mode.

## QUANTIFICATION AND STATISTICAL ANALYSIS

### Single-cell methylation and multi-omics data mapping (alignment, quality control (QC))

The snmC-seq3, snmCT-seq and snm3C-seq mapping were using the YAP pipeline (cemba-data package, v1.6.8, https://hq-1.gitbook.io/mc/) as previously described.^8,19^ The major steps of the processing steps include:

1) Demultiplexing FASTQ files into single cells (cutadapt^70^, v2.10);
2) reads level QC;

For snmCT-seq (methylome part):

3a) Reads from step 2 were mapped onto human hg38 genome (one-pass mapping for snmCT-seq, two-pass mapping for snm3C) (bismark^71^ v0.20, bowtie2^72^, v2.3);

4a) PCR duplicates were removed using Picard MarkDuplicates, the non-redundant reads were filtered by MAPQ > 10. To select genomic reads from the filtered BAM, we used the ‘‘XM-tag’’ generated by Bismark to calculate reads methylation level and keep reads with mCH ratio < 0.5 and the number of cytosines ≽ 3.

5a) Tab-delimited (ALLC) files containing methylation level for every cytosine position were generated using allcools ^69^ (v1.0.8) bam-to-allc function on the BAM file from step 4a.

For snmCT-seq (RNA part):

3b) To map transcriptome reads, reads from step 2 were mapped to GENCODE human v30 indexed hg38 genome using STAR^73^ (v2.7.3a) with the following parameters: --alignEndsType Local --outSAMstrandField intronMotif --outSAMtype BAM Unsorted --outSAMunmapped None --outSAMattributes NH HI AS NM MD --sjdbOverhang 100 --outFilterType BySJout --outFilterMultimapNmax 20 --alignSJoverhangMin 8 --alignSJDBoverhangMin 1 --outFilterMismatchNmax 999’ # ENCODE standard options –outFilterMismatchNoverLmax 0.04 --alignIntronMin 20 --alignIntronMax 1000000 --alignMatesGapMax 1000000 --outFileNamePrefix rna_bam/TotalRNA

4b) The STAR mapped reads were first filtered by MAPQ > 10. To select RNA reads from the filtered BAM, we used the ‘‘MD’’ tag to calculate reads methylation level and kept reads with mCH ratio > 0.9 and the number of cytosines ≽3. The stringency of read partitioning was tested previously.^8^

5b) BAM files from step 4b were counted across gene annotations using featureCount^74^ (1.6.4) with the default parameters. Gene expression was quantified using either only exonic reads with ‘‘-t exon’’ or both exonic and intronic reads with ‘‘-t gene.’’

For snm3C-seq (3C modality part):

4b) After the initial mC reads alignment as above, unmapped reads were retained and splitted into 3 pieces by 40bp, 42bp, and 40bp resulting in six subreads (read1 and read2). The subreads derived from unmapped reads were mapped separately using HISAT-3N ^75^ adapted in YAP pipeline (cemba-data package). All aligned reads were merged into BAM using Picard SortSam tool with query names sorted. For each fragment, the outermost aligned reads were chosen for the chromatin conformation map generation. The chromatin contacts and following analysis were processed using the scHiCluster described previously.^76^ (https://zhoujt1994.github.io/scHiCluster/intro.html)

### Preprocessing of snmC-seq, snmCT-seq and snm3C-seq data

Primary QC for DNA methylome cells was (1) overall mCCC level < 0.05; (2) overall mCH level < 0.2; (3) overall mCG level < 0.5; (4) total final DNA reads > 100,000 and < 10,000,000; and (5) Bismarck mapping rate > 0.5. Note that the mCCC level estimates the upper bound of the cell-level bisulfite non-conversion rate. Additionally, we calculated lambda DNA spike-in methylation levels to estimate each sample’s non-conversion rate. For the transcriptome modality in snmCT-seq, we only kept the cells containing < 5% mitochondrial reads, total RNA reads > 5,000. For snm3C-seq cells, we also required cis-long-range (two anchors > 2500 bp apart) > 50,000.

### Clustering analysis of snmCT-seq and snm3C-seq data

For snmCT-seq (RNA part):

The whole gene RNA read count matrix was used for snmCT-seq transcriptome analysis. Cells were filtered by the number of genes expressed > 1,000 and genes were filtered by the number of cells expressed > 10. The count matrix X was then normalized per cell and transformed by ln(X + 1). After log transformation, we used the scanpy.pp.highly_variable_genes to select the top genes based on normalized dispersion. The selected feature matrix was scaled to unit variance and zero mean per feature followed by PCA calculation. To correct batch effects across individuals, we established a highly efficient framework based on the Seurat R integration algorithm.^77^ The integration framework consisted of 3 major steps to align snmCT-seq datasets on fibroblasts and iNs from different donors onto the same space: (1) using dimension reduction to derive embedding of the multiple datasets separated by donors in the same space; (2) using canonical correlation analysis (CCA) to capture the shared variance across cells between datasets and find anchors as 5 mutual nearest neighbors (MNN) between each two paired datasets; and (3) aligning the low-dimensional representation of the paired data sets together with the anchors.

To consensus clustering based on fixed resolution parameters (range from 0.2 to 0.6), we first performed Leiden clustering ^78^ 200 times, using different random seeds. We then combined these result labels to establish preliminary cluster labels. Following this, we trained predictive models in the principal component (PC) space to predict labels and compute the confusion matrix. Finally, we merged clusters with high similarity to minimize confusion. The cluster selection was guided by the R1 and R2 normalization applied to the confusion matrix, as outlined in the SCCAF^79^ package. This framework was incorporated in “ALLCools.clustering.ConsensusClustering” function.

For snmCT-seq (methylome part):

We performed clustering analysis with the mCH and mCG fractions of chrom100k matrices described previously.^8^ Most functions were derived from allcools^69^, scanpy^80^ and scikit-learn^81^ packages. In general, the major steps in the clustering included: (1) feature filtering based on coverage, exclude ENCODE blacklist and located in autosomes; (2) Highly Variable Feature (HVF) selection; (3) generation of posterior chrom100k mCH and mCG fraction matrices; (4) clustering with HVF and calculating Cluster Enriched Features (CEF) of the HVF clusters with “ALLCools.clustering.cluster_enriched_features” function; (5) calculating PC in the selected cell-by-CEF matrices and generating the t-SNE^82^ and UMAP^83^ embeddings for visualization; and (6) consensus clustering process using “ALLCools.clustering.ConsensusClustering” function.

### Identification of DEGs, DMRs and aDMRs enriched hotspots

After finalizing clustering in *in vitro* fibroblast-iN snmCT-seq data analysis, we used a paired strategy to calculate RNA DEGs within a specific cluster for AD-specific (AD versus CTRL) or within a specific individual line for conversional DEGs (fibroblast versus iNs). We used all the protein-coding and long non-coding RNA genes from hg38 gencode v30 with the scanpy.tl.rank_genes_group function with the Wilcoxon test and filtered the resulting marker gene by adjusted P value < 0.01 and log2(fold-change) > 1.

For DMRs identification, we merged the single-cell ALLC files into pseudo-bulk level using the “allcools merge-allc” command. Next, we performed DMR calling with methylpy^84^ on a grouped pseudo-bulk allc files. For example, to identify AD-specific methylation signatures in fibroblasts and iN clusters, we merged the samples from all individuals in AD and CTRL groups separately and then we called DMRs between these two groups. After getting the primary set of DMRs, we counted the methylation level at these DMRs from all individuals using the “methylpy add-methylation-level” function. Additional filtering on the DMRs was performed by comparing the methylation levels among different individuals within groups using Student’s t-test. Only DMRs with a minimum p-value less than 0.05 between any two groups were retained. The same processes were used to identify aDMRs in each specific brain cell type of snm3C-seq datasets. aDMRs enriched hotspots of the *in vivo* entorhinal cortex were identified by a sliding window of 5kb bin across the autosomes, with normalized GC content. We employed PyComplexHeatmap ^85^ to visualize methylation level at these DMRs in the complex heatmaps. Hypomethylated DMRs in the corresponding sample groups and cell types were labeled for better visualization. The heatmap rows were split according to sample groups, and the columns were split based on DMR groups and cell types. Within each subgroup, rows and columns were clustered using ward linkage and the Jaccard metric. The aDMRs-enriched hotspots were visualized by tagore package.^86^

### Gene set enrichment test, motif enrichment, chromatin states and functional enrichment of DMRs

To validate the DEGs found in snmCT-seq dataset *in vitro* fibroblast/iN models, we performed GO enrichment test using GSEApy ^87^ and Enrichr ^88^ open source. The −log(adjusted P value) of KEGG pathway enrichment in each selected gene set was color-coded on the enrichr combined score with KEGG terms. For motif enrichment analysis, we obtained the hypomethylated and hypermethylated DMRs reported by methylpy from the columns ‘hypermethylated_samples’ and ‘hypomethylated_samples’. HOMER was used to identify enriched motifs within these different sets of DMRs for each comparison. The results from HOMER’s ‘knownResults.txt’ output files were used for downstream analysis. Only motif enrichments with a p-value < 0.01 were retained. The motif enrichment results were visualized using scatterp‘ots in seaborn. To perf’rm fu‘ctional enrichment analysis of DMRs, we utilized GREAT (http://great.stanford.edu/public/html/index.php). The genome feature annotation of aDMRs enriched hotspots and ML identified DMSs in the entorhinal cortex was conducted using “annotatePeaks.pl” functions in HOMER. The chromHMM states enrichment analysis of aDMRs were quantified by “bedtools intersect” the overlapping of aDMRs with the corresponding ChromHMM states based on histone ChiP-Seq peaks from the Roadmap Epigenomics project derived from frontal cortex (67 and 80 years old female donors), the accession number is ENCSR867UKF in the ENCODE database. Enrichment tests were performed using Fisher’s tests with the significance of FDR adjusted p-value calculated by multiple tests.

### Integration and annotation between snm3C-seq datasets and human brain atlas

To integrate our snm3C-seq dataset to the reference human brain methylation atlas (HBA),^6^ we used methylation information from both CHN and CGN sites. We derived log scaled cell-by-100kb-bin methylation fraction matrices for CGN and CHN separately. After removing all low quality bins (hg38 genome blacklist, coverage<500, or coverage>3000), we selected features that were both highly variable and cluster enriched in HBA for PCA. The first 100 PCs of mCG and mCH matrices were normalized by their standard deviations and then concatenated horizontally for integration. We used canonical correlation analysis (CCA) to capture the shared variance across cells between datasets and then selected 5 mutual nearest neighbors (MNNs) as anchors between the datasets. Next, we used HBA as a reference dataset to pull our dataset into the same space. More details on our integration algorithms are described in Tian et al.^6^ Lastly, we used Harmony on the CCA integrated matrix for better integration between individuals. After integration, we annotated major cell types by the most numerous HBA cell type within each leiden cluster in the joint embedding.

### Chromation contact matrix and preprocessing imputation of snm3C-seq datasets

We performed imputation using scHiCluster (https://zhoujt1994.github.io/scHiCluster/intro.html) to the contact matrices at 100kb, 25kb and 10kb resolution for single cell contacts within 10.05Mb (100kb and 25kb), and 5.05Mb (10kb). For imputation at 10kb resolution specifically, convolution and random walk were performed to speed up the imputation. For pseudo-bulk analysis, we merged cells from each donor by major cell type (ASC, MGC, ODC, Inh, Ex) with cell number across individuals as closely as possible to reduce bias created by different sequencing depth. Most cell types across individuals had at least 150 cells for pseudo-bulk analysis. For pseudo-bulk analysis that compared AD and CTRL cell types, the same number of cells (n=400) were randomly selected and merged among AD and CTRL individuals.

### Contacts, loop, domain, and compartment analysis

As described above, pseudo-bulk cell type groups were merged by individual and disease status. We used imputed contact matrices for both single-cell and pseudo-bulk domain calling at 25 kb resolution and loop calling at 10 kb resolution. Raw contact matrices were used instead to infer A/B compartments for pseudo-bulk groups at 100k resolution to better capture detailed genome interaction. Differential loops, domains, and compartments were derived as described previously,^6^ so as saddle plots, compartment strengths, and loop summits. We calculated the cis (intra-chromosomal) contact probability normalized by CG counts for each cell. DNA contacts were binned by an exponent step of 0.125 with a base of 2, ranging from contact distance between 2500bp to 249Mb. The start and end of the bin were calculated by 2500×2^0. 125*i* and 2500×2^0. 125(*i* + 1). The short-long ratio in Fig.4C was defined as the mean probability of contact in 51st (200k) to 76th (2M) binds divided by the probability of contact in 103rd (20M) to 114th (50M) bins. Based on the loop interactions mapped in 3C contacts, we assigned aDMRs to a gene if the transcription start site (TSS) located in one 10kb-bin had interactions with the bin where aDMRs were located. Then, aDRMs were considered as putative CREs of DEGs if the aDMRs paired with genes found differential expressed in published sn-RNA datasets between AD and CTRL.^89^

### Determination of reliable CpG sites for AD prediction

Identifying aDMRs to predict AD is multifaceted, involving various steps from preprocessing and feature selection to validation.

#### Data preprocessing

Our initial step was to merge DMR sites between AD and CTRL groups for every cell type. The methylation fraction was then extracted for all these sites for every sample. To maintain data reliability, we filtered out any sites where the change in the methylation fraction across samples was less than 0.4, or the standard deviation was less than 0.1. Given the inherent biases in sample data, we further normalized these data within each sample using the z-score. Following this preprocessing, the resultant data served as the primary candidate set we considered for subsequent feature selection.

#### Feature selection

We employed an iterative feature selection approach to ensure a comprehensive feature selection that captured as much reliable and informative data as possible. This was done over 30 rounds. We used stratified 3-fold cross-validation (CV) in every round to train Random Forest classifiers (RFCs). The importance of the remaining features was gauged by the average feature importance derived from the RFCs. The top 500 features were chosen in every round, and the rest were reserved for the next round. The parameters set for the RFCs included utilizing 500 trees with a max_depth of 3 for each RFC.

#### Method evaluation

To ascertain the predictive capability of our selected features, we performed a stratified 4-fold CV, ensuring that the stratification was based on the combined label of AD versus CTRL and *in vivo* versus *in vitro* conditions. In each fold, the 3 training subsets underwent the feature selection process mentioned earlier. Following this, an RFC was trained based on the chosen features, which was then used on the remaining fold to determine its accuracy. After completing the 4-fold CV, we found that the overall prediction accuracy was 97.1%.

#### Mitigating donor effects

It is essential to account for donor variability. To do this, we selected shared features from the prior 4-fold CV as candidates. We then evaluated their importance in predicting AD vs. CTRL or determining the donor. Figure 7E demonstrates how each feature played a role in these 2 prediction tasks. We finalized only those features that held a positive importance for AD vs. CTRL predictions and an importance of less than 5e-4 for donor predictions. This meticulous process resulted in 859 CpG sites.

#### Final predictor and validation

To validate our method, we trained an RFC using the 859 selected sites and then applied it to a separate snmC dataset comprising individual repeats and 3 unseen donors. This resulted in an accuracy of 100%.

## ACKNOWLEDGMENTS

We thank the brain/fibroblast tissue donors and their families; this work would not be possible without them. This research was supported by an AHA-Allen Initiative in Brain Health and Cognitive Impairment award made jointly through the American Heart Association and The Paul G. Allen Frontiers Group: 19PABH134610000. This work was supported by the Stem Cell Core Facility of the Salk Institute with funding from the Helmsley Charitable Trust, the Shiley-Marcos Alzheimer’s Disease Research Center (ADRC; AG062429) at the University of California, San Diego (UCSD), and the National Institute of Aging of the National Institutes of Health (P30AG068635). We thank the members of the AHA-Allen community and EAB members for the discussion. We thank Dr. Stefy Zambetti for outstanding administrative assistance for the AHA-Allen initiative at Salk. This work utilized the Stampede2 supercomputing resources at the Texas Advanced Computing Center through allocation MCB130189 from the Extreme Science and Engineering Discovery Environment. We are very grateful to members of the Ecker group for their feedback and discussion.

## AUTHOR CONTRIBUTIONS

J.R.E., F.H.G., B.W., and J.R.J. conceived the study. J.R.E. and F.H.G. supervised the study. C.L. and B.W. developed the snmCT-seq method. B.W., A.B., R.C., J.R.N., M. K., J. A., C.V., M.L., N.C., and C.O. generated the snm3C-seq, snmCT-seq and snmC-seq data. B.W., J.Z., W.T., Y.W., W.W. analyzed snm3C-seq, snmCT-seq and snmC-seq data. J. M. acquired human brain specimens. J.R.J., A. K., and R. G. generated the iN cells. B.W., J.Z., and W.T. drafted the manuscript. J.R.E., F.H.G., and J.R.J. edited the manuscript.

## Supplementary Figures

Supplementary Figure S1. QC metrics of snm3C-seq in entorhinal cortex and cell type-specific aDMRs.

Supplementary Figure S2. DEGs linked by loop interaction with aDMRs.

Supplementary Figure S3. Chromosomal epigenome erosion occurs in multiple cell types.

Supplementary Figure S4. Fibroblast iNs retain aging-related methylation signatures compared to iPSCs derived iNs.

Supplementary Figure S5. snmCT-seq quality control (QC) metrics and cell state annotation.

Supplementary Figure S6. Motif enrichment analysis of aDMRs and their linked DEGs in in vitro cellular models.

Supplementary Figure S7. Examples of ML selected features in AD predictors.

## Supplementary Tables

Supplementary Table S1. Donor information

Supplementary Table S2. aDMRs enriched hotspot in entorhinal cortex

Supplementary Table S3. aDMRs linked DEGs by loop interactions in brain cell types

Supplementary Table S4. Cell state marker genes of *in vitro* fibroblast direct converted iN

Supplementary Table S5. DEGs in Fib/iN and GO analysis gene terms

Supplementary Table S6. Shared aDMRs between *in vitro* and *in vivo*,and ML-selected CpG sites

---

**Figure S1:**
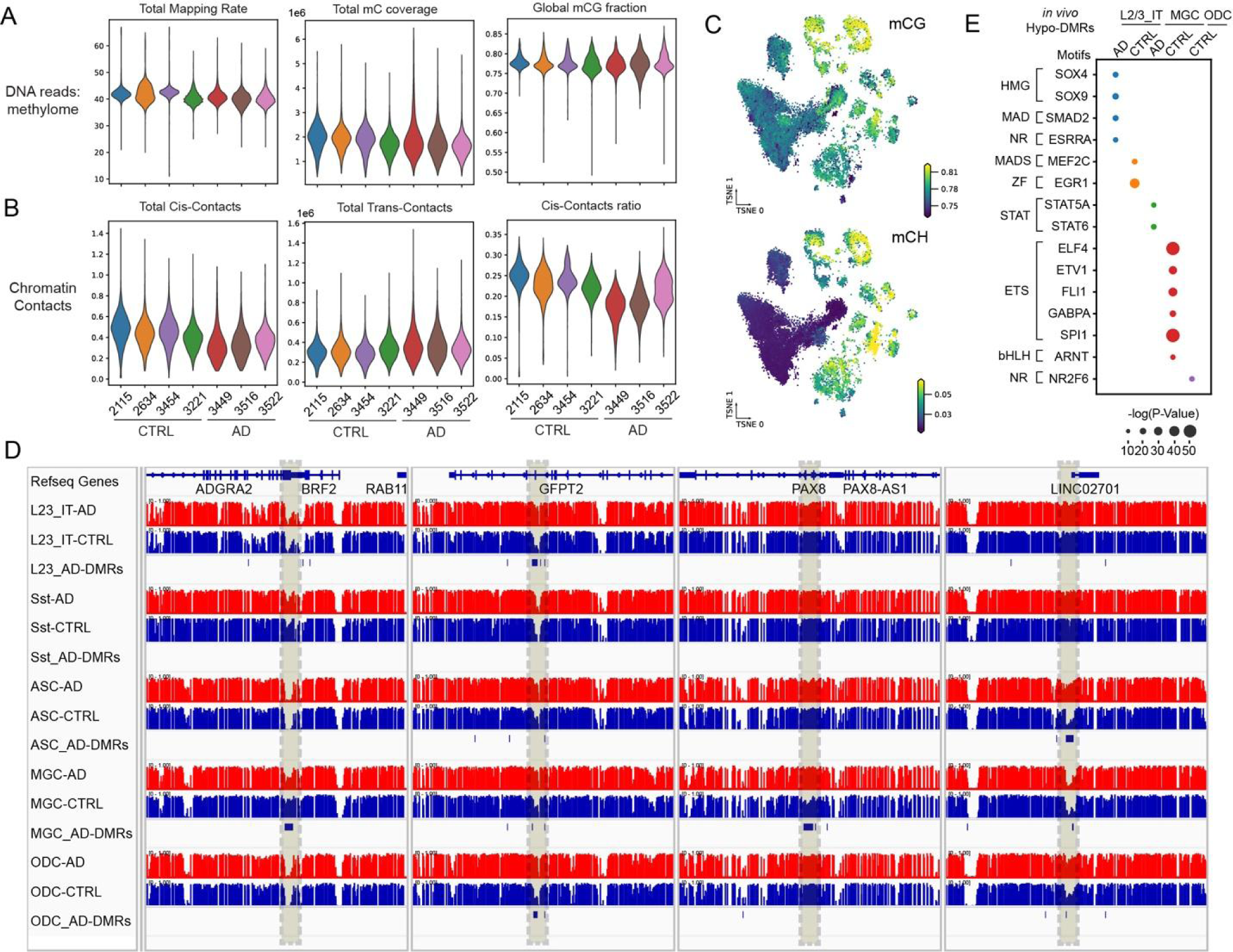
QC metrics of snm3C-seq in entorhinal cortex and cell type-specific aDMRs. (A) DNA modality QC metrics per cell grouped by individuals on distribution of overall mapping rate, total reads number and global mCG level. (B) Chromatin conformation modality QC metrics per cell grouped by individuals on distribution of total cis-contact reads, total trans-contact reads and cis–contact ratio. (C) t-SNE visualization of global methylation level at CG, CH and CCC context. (D) Examples of cell type-specific aDMRs. (E) Scatter plot shows the significant enrichment of motifs (E, −log(p-value) > 15) at hypomethylated aDMRs identified in corresponding brain cell types, with the size of the dot indicating the −log(p-value).

**Figure S2:**
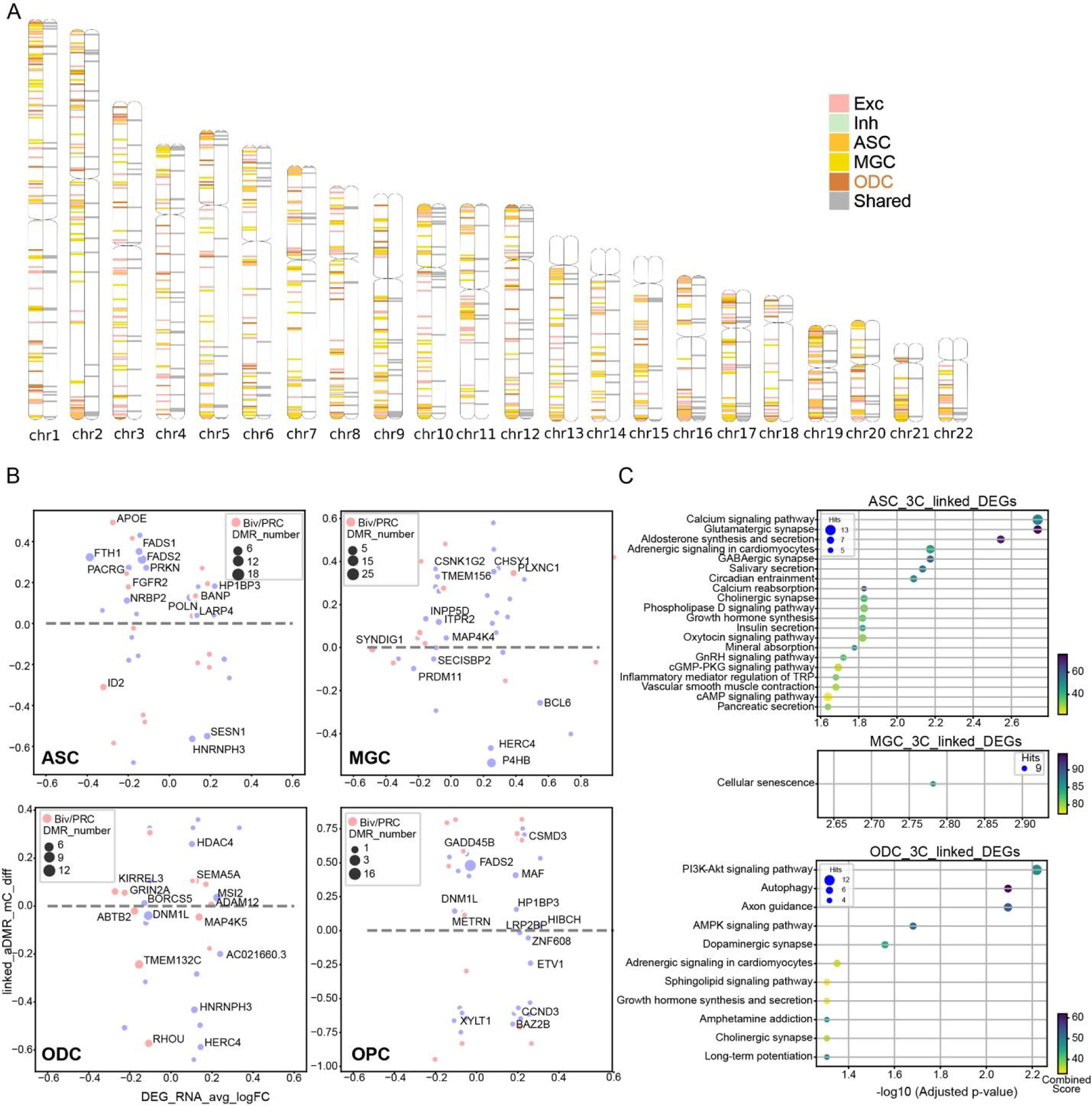
DEGs linked by loop interaction with aDMRs. (A) 1,795 hotspots of aDMRs across the total autosomes. 5kb-bin hotspots are flanked to 1M only for visualization and colored by cell types where hotspots are identified from. The left chromosome copy shows the cell type specific hotspots, the right copy shows the hotspots found in at least two brain cell types. (B) Scatter plot of the average log2 RNA expression fold changes (AD/CTRL) of DEGs as X-axis and average methylation difference (AD-CTRL) of DEGs linked aDMRs in glia cell types (ASC, MGC, ODC and OPC). The size and color represent the number of linked aDMRs and whether the promoter of the gene is on repressive chromHMM states (TssBiv and ReprPC). (C) KEGG pathway enrichment analysis for aDMRs linked DEGs in ASC, MGC, ODC and OPC. X-axis shows the enrichment significance as −log (adjusted p-value), and Y-axis represents pathways. The size and color represent the number of related genes and combined score, respectively.

**Figure S3:**
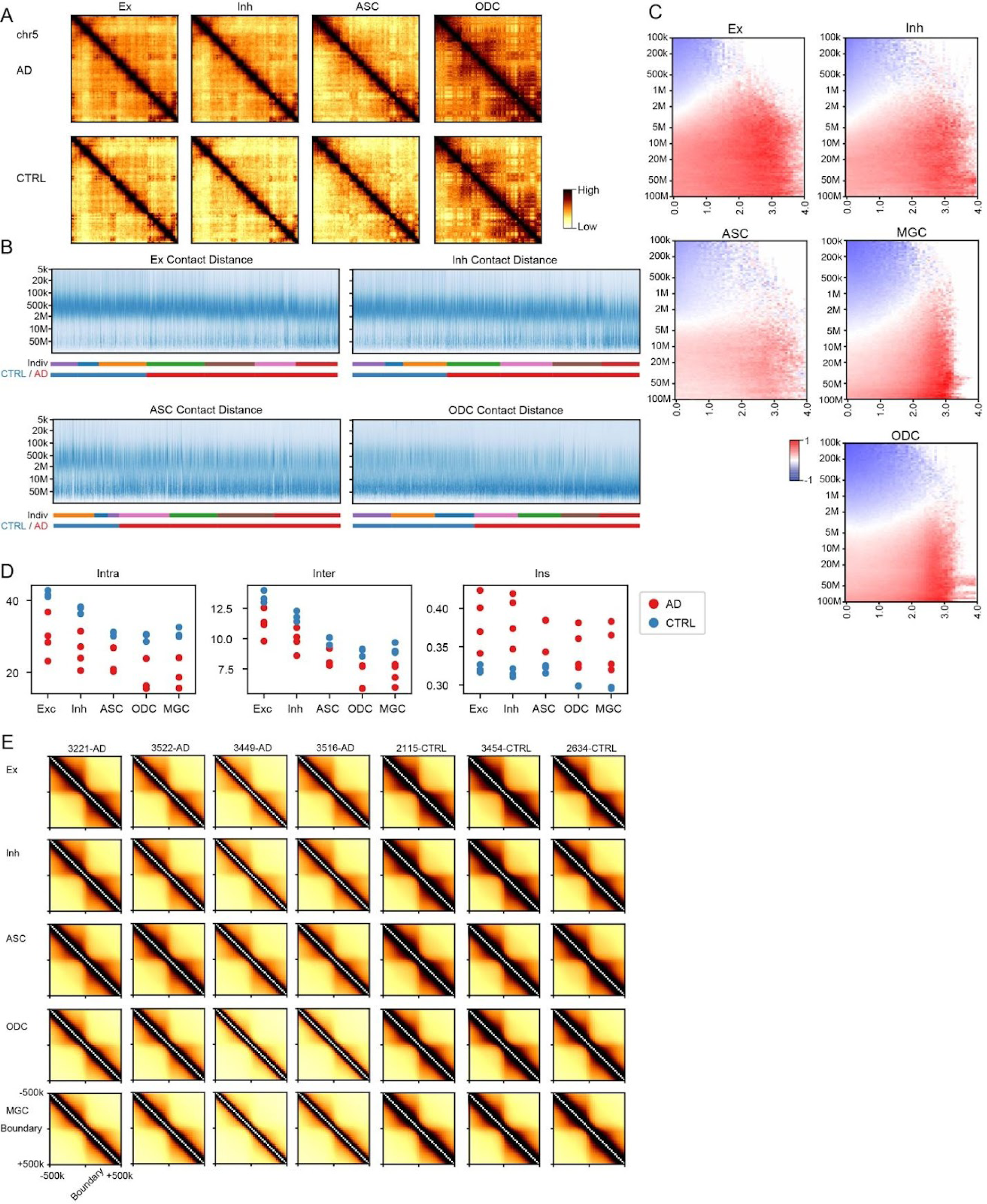
Chromosomal epigenome erosion occurs in multiple cell types. (A) Chromatin contact map of excitatory neurons (Ex), inhibitory neurons (Inh), astrocytes (ASC) and oligodendrocytes (ODC) from AD and CTRL across chromosomes, chr5 as an example. (B) Frequency of contacts against genomic distance in each single cell of Ex, Inh, ASC and ODC cell types, Z-score normalized within each cell (column). The y-axis is binned at log2 scale. Each cell in the x-axis is grouped and colored by individuals and CTRL or AD. (C) The log2 ratio between AD and CTRL of frequency of contacts grouped by genomic distance and the difference of raw compartment scores at the two anchors of a contact across cell types. (D) Contact scores between the intra-compartment, inter-compartment regions and the ratio between inter- and intra-compartment interactions. (E) Contacts correlation around domain boundaries of major types across individuals.

**Figure S4.**
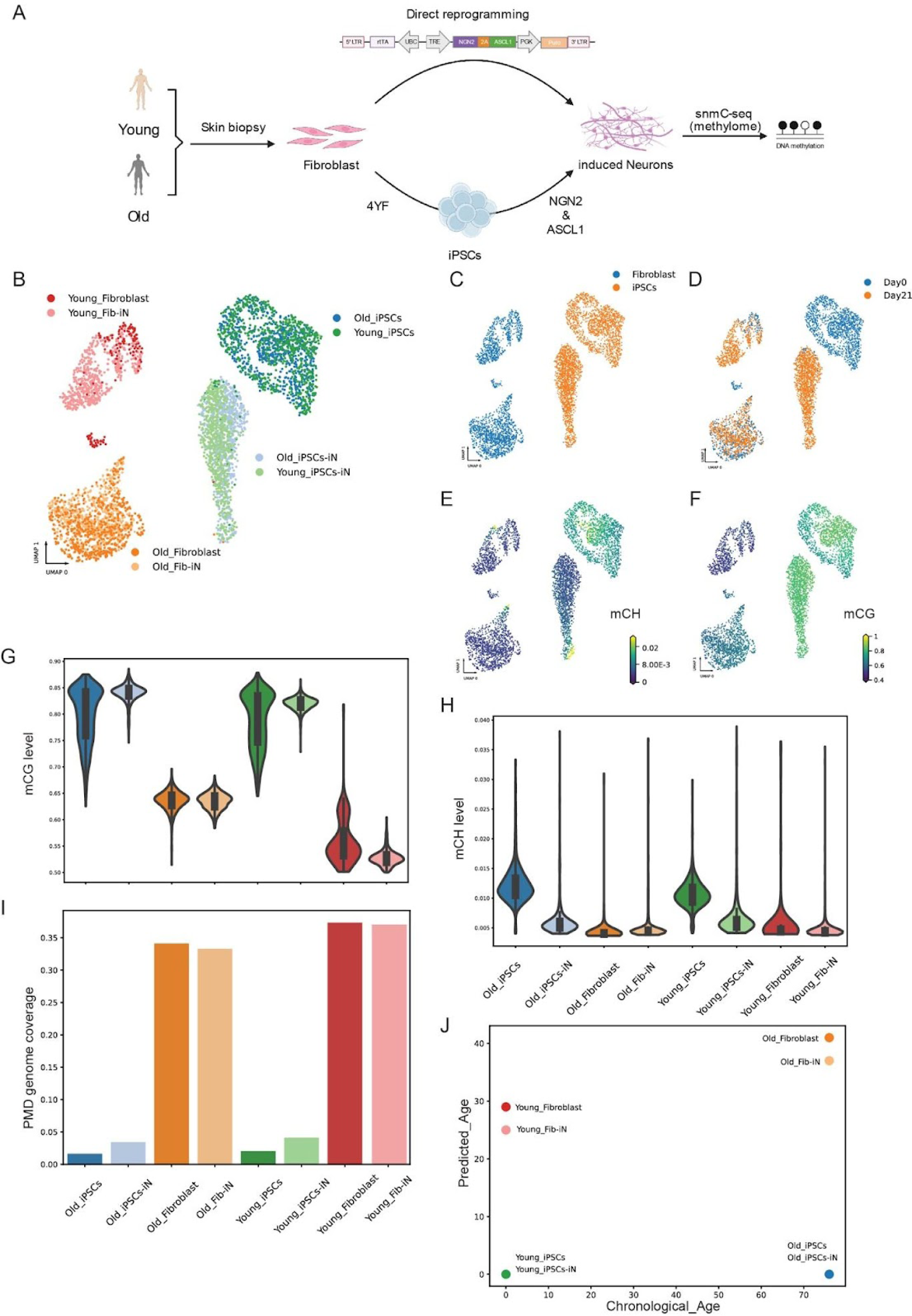
Fibroblast iNs retain aging-related methylation signatures compared to iPSCs derived iNs. (A) Schematic representation of comparing iPSC-differentiated iNs and fibroblast directly converted iNs via inducible expression of the same proneuronal factors NGN2-2A-ASCL1 (N2A). (B-D) Clustering and annotation of iNs generated via two methods based on methylation levels of 100kb bins. Cells were colored by samples (B), starter cell sources (C), and collection days (D). (E-F) Global methylation levels of mCH (E) and mCG (F) on 2D umap clustering. (G-H) Violin plots showing the average levels of mCG (G) and mCH (H) across donor and cell types. (I) The genome coverage consists of partially methylated domains (PMDs) across cell types. (J) DNAm age prediction of fibroblast/iNs and iPSC/iNs by multi-tissue age estimator ^90^ compared to chronological age.

**Figure S5.**
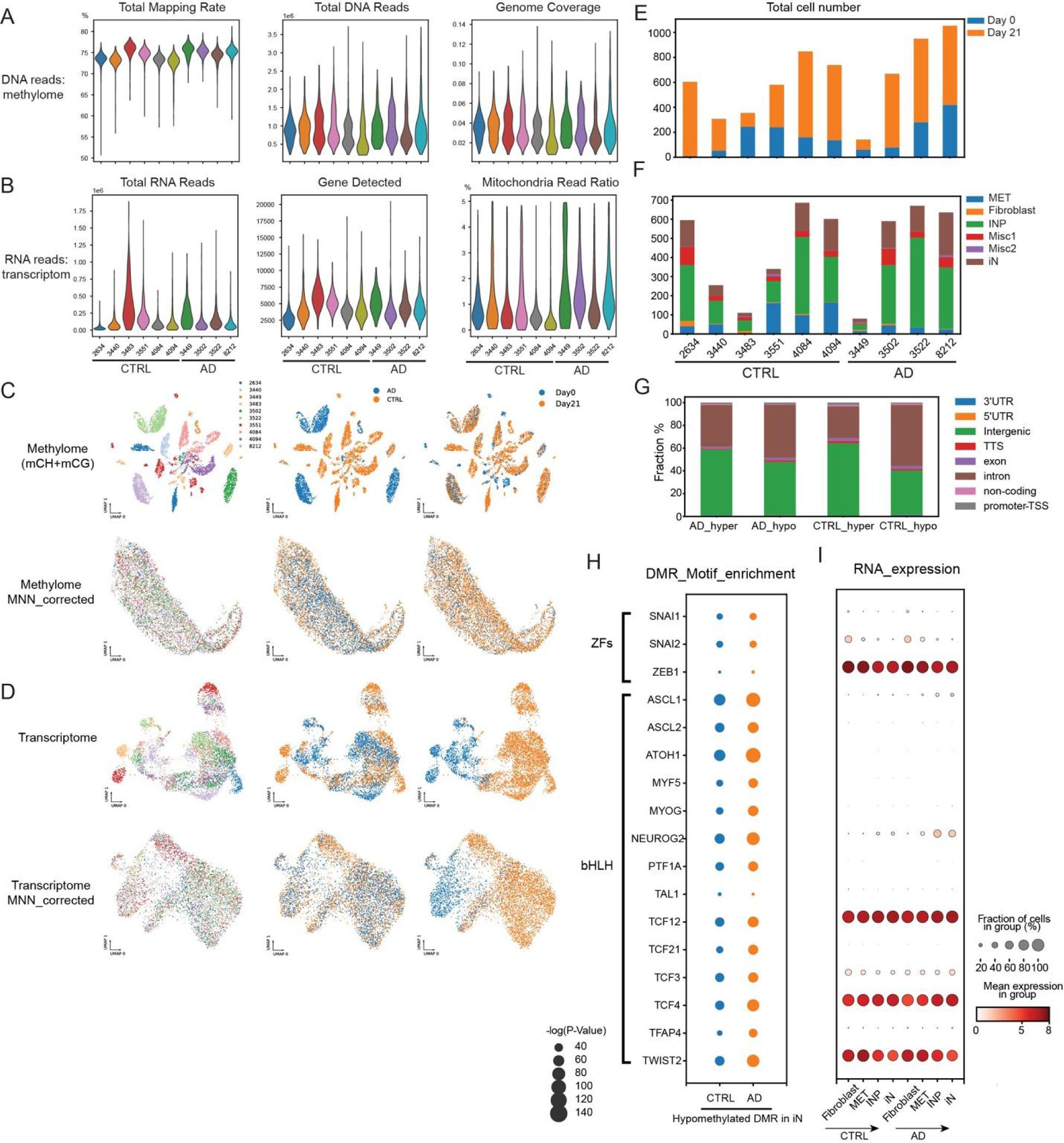
snmCT-seq quality control (QC) metrics and cell state annotation. (A) DNA modality QC metrics per cell grouped by individuals on distribution of total mapping rate, total reads number and genome coverages. (B) RNA modality QC metrics per cell grouped by individuals on distribution of total RNA reads, gene detected, and mitochondria read ratio. (C) UMAP embedding before and after batch correction of mCG and mCH level at 100kb bins. Cells were colored by individuals, groups and collection day separately. (D) UMAP embedding before and after batch correction of RNA expression. Cells were colored by individuals, groups and collection day separately. (E-F) Cell number distribution of collection day (E) and cell states on Day 21 (F) across individuals. (G) Genomic feature fraction of conversional DMRs grouped by hypermethylated or hypomethylated between fibroblasts versus iNs in AD and CTRL groups. (H-I) Scatter plot shows the enrichment of TF motifs (H) at hypomethylated conversional DMRs in iNs compared to fibroblasts in AD and CTRL, with the size of the dot indicating the −log(p-value). The corresponding RNA expression of TF candidates is shown in (I), the size of dot presents the fraction of expressed cells whereas colors show the mean expression (log (CPM+1)) of cell populations.

**Figure S6:**
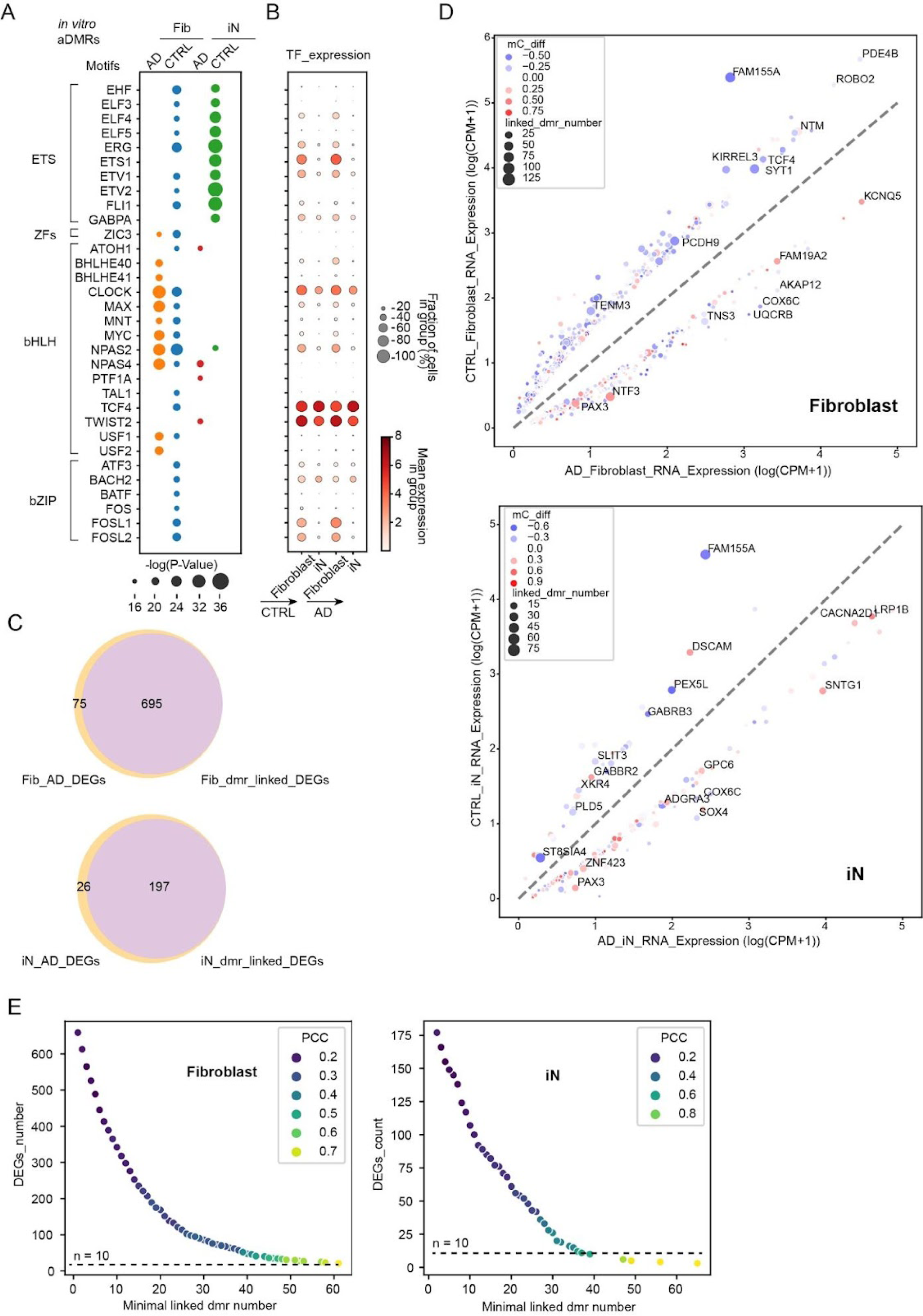
Motif enrichment analysis of aDMRs and their linked DEGs in *in vitro* cellular models. (A-B) Scatter plot shows the motif enrichment analysis of the hypomethylated aDMRs in AD and CTRL in fibroblasts/iNs states, with the size of the dot indicating the −log(p-value). The corresponding RNA expression of TF candidates is shown in (B), the size of dot presents the fraction of expressed cells whereas colors show the mean expression (log (CPM+1)) of cell populations. (C) Venn diagrams showing the overlapping of shared DEGs identified in fibroblast with DEGs linked with aDMRs by GREAT algorithm.^63^ (D) RNA expression scatter of DEGs in fibroblasts (upper panel) and iN (lower panel) with linked aDMRs by GREAT algorithm. X axis and Y axis present log2 value of normalized gene expression as CPM (Counts per million) in AD and CTRL samples. The size and color represent the number of linked aDMRs and average methylation changes, respectively. Top DEGs’ names are labeled beside the dot. (E) The distribution of DEGs number in fibroblast and iN over the minimal number of aDMRs associated by GREAT algorithm, colored by the Pearson correlation coefficient between log2 fold changes of RNA expression (AD/CTRL) and average methylation difference (AD-CTRL) of associated aDMRs by GREAT algorithm.^63^

**Figure S7:**
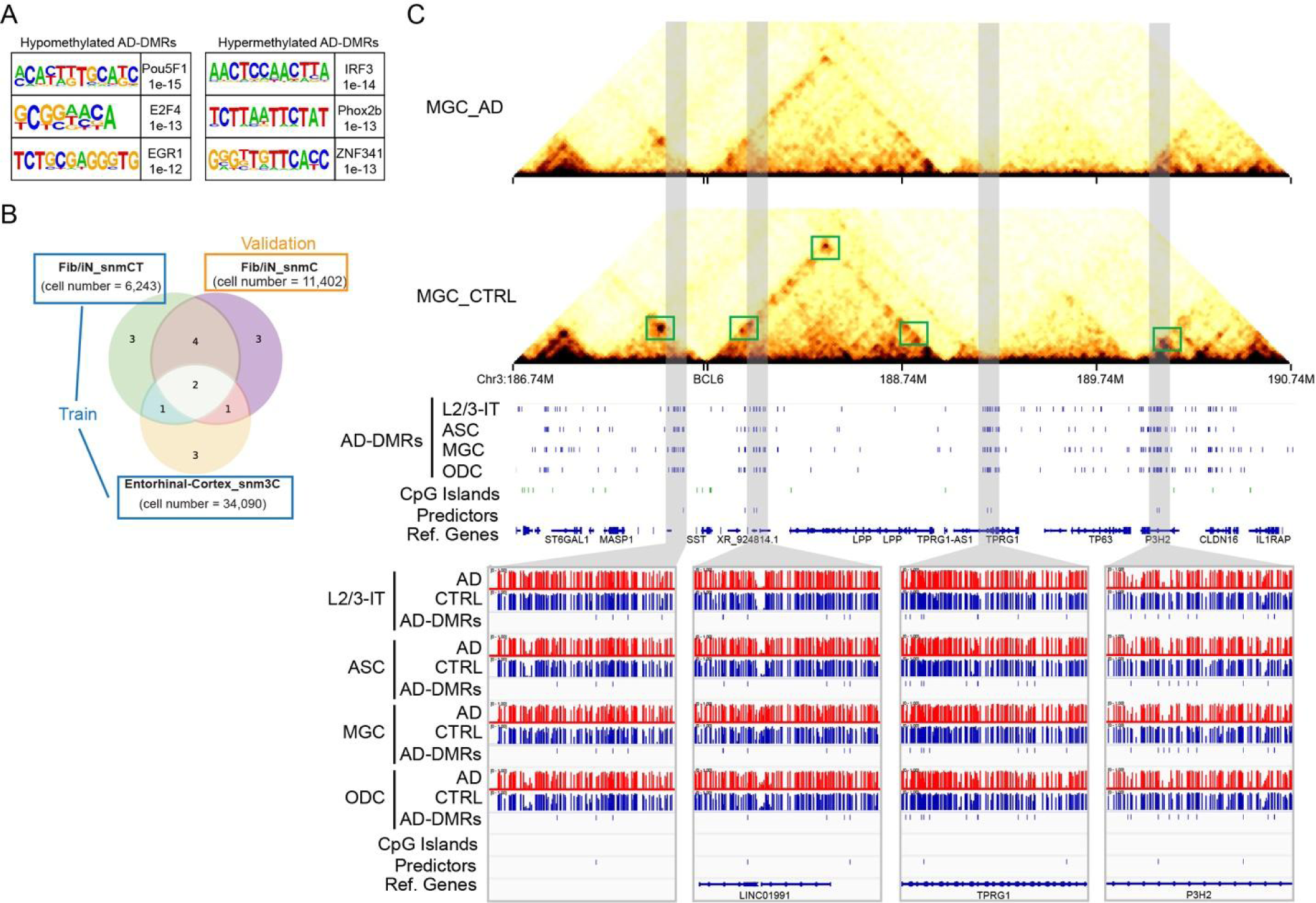
Examples of ML selected features in AD predictors. (A) Top 3 motif enrichment of shared aDMRs (B) Datasets used for training and testing of AD predictors. (C) De novo selective AD predictors by machine learning are overlapped with shared aDMRs across cell types. Upper panel, chromatin conformation around the gene BCL6 shows the decreased loop contacts in AD microglia. Middle panels, the selected features in the AD predictor are located at LINC01991, TPRG1 and P3H2 intronic regions, overlapped with shared aDMRs across cell types. Lower panel, browser view of methylation level surrounds the AD predictors in AD and CTRL.

